# The molecular properties of the bHLH TCF4 protein as an intrinsically disordered hub transcription factor

**DOI:** 10.1101/2025.02.06.636797

**Authors:** Sozańska Nikola, Barbara P. Klepka, Niedźwiecka Anna, Żukowa Lilia, Dadlez Michał, Greb-Markiewicz Beata, Ożyhar Andrzej, Tarczewska Aneta

## Abstract

1.

**Background:** Transcription factor 4 (TCF4) is a member of the basic helix-loop-helix (bHLH) family of transcription factors and guides proper embryogenesis, particularly neurogenesis, myogenesis, heart development and hematopoiesis. bHLH proteins exhibit variations in length, expression patterns and the presence of defined motifs. The interaction of TCF4 with DNA is dependent on the presence of a conserved bHLH domain, particularly the presence of a basic (b) motif. Most mutations in the *Tcf4* gene are either associated with the development of serious nervous system disorders, such as Pitt-Hopkins syndrome or schizophrenia, or are lethal. Although TCF4 is essential for the proper development and function of the human body, apart from the isolated bHLH domain, there is a lack of fundamental knowledge about the structure of TCF4.

**Methods:** Recombinant full-length TCF4 was expressed in *E. coli* cells and purified using chromatographic techniques. To determine the conformation of the polypeptide chain of TCF4, we performed *in vitro* analysis using hydrogen/deuterium exchange mass spectrometry (HDX-MS). We then determined the dissociation constant (K_D_) of the TCF4:DNA complex using independent methods, including fluorescence polarization (FP), electrophoretic mobility shift assay (EMSA), and fluorescence correlation spectroscopy (FCS). Finally, we compared the properties of TCF4 in its *apo* and *holo* form in relation to the changes of the conformation of the polypeptide chain (HDX-MS), hydrodynamic properties (e.g., sedimentation-velocity analytical ultracentrifugation; SV-AUC), and stability (thermal shift, circular dichroism; CD).

**Results:** In this study, we demonstrate the molecular characteristics of full-length TCF4, the dimer of which is one of the largest intrinsically disordered proteins (IDPs) described to date. According to our findings, the structure of TCF4 is extensively disordered. Only the bHLH domain, which comprises 10% of the polypeptide chain, exhibits a stable fold. Strikingly, Ephrussi-box (E-box) binding via the bHLH domain has no significant effect on the disordered nature of TCF4, but it does influence the stability of TCF4.

**Conclusions:** We suggest that bHLH plays the role of an anchor localizing TCF4 to specific gene sequences. The dual nature of the TCF4 structure, as well as the fact that the intrinsically disordered regions (IDRs) represents most of the protein sequence, suggests that TCF4 may act as a hub transcription factor regulating the expression of specific genes through the interaction of IDRs with gene-specific partners.

## 3. Background

Transcription factor 4 (TCF4) is a mammalian transcription factor (TF) also known as immunoglobulin transcription factor 2 (ITF2) or SL3-3 enhancer factor 2 (SEF2). TCF4 is expressed in several tissue types, but mainly in the nervous system, where the protein is essential for brain development, memory and cognition. Mutations in the *Tcf4* gene have been associated with schizophrenia, autism spectrum disorder and Pitt-Hopkins syndrome [1]. TCF4 is a member of the class I bHLH (basic helix-loop-helix) family [2]. Class I bHLH proteins, called E proteins, are related to *Drosophila melanogaster* Daughterless, whereas the mammalian members are TCF4, TCF3 and TCF12 [2,3]. The human *Tcf4* gene (ID 6925) is located on chromosome 18 in the region 18q21.2 and it spans 437 kb. It consists of 41 exons, 21 of which are alternative 5’ coding exons. These exons, together with open reading frames and possible translation start codons, are responsible for the existence of many isoforms of TCF4 (A– R). Isoforms with unique N-terminal sequences are generated via the use and splicing of alternative 5’ exons [4].

TCF4 regulates gene transcription in a ligand-independent manner. It was shown to act as a repressor [5–7] or activator [8–10] of transcription depending on the interacting partner in the heterodimer. TCF4, as well as other class I bHLH proteins, can interact and heterodimerize with class II members, such as achaete-scute homolog 1 (ASCL1) or atonal bHLH transcription factor 1 (ATOH1), which are potent inducers of cell type specification [11,12]. The dimerization partners of TCF4 also include intellectual disability (ID) proteins from class V and hairy and enhancer of split (HES) proteins from class VI of the bHLH family. The interaction of TCF4 with the DNA-binding protein inhibitor ID-2 (ID-2) results in the formation of an inactive heterodimer that binds DNA but is unable to bind activators that are essential for the activation of transcription [13]. The transcriptional activity of class I bHLH proteins is also known to be repressed upon binding/heterodimerization with HES proteins [14,15].

The first localization studies of human TCF4 reported the presence of the protein in the cell nucleus [16]. Later, exclusive localization in the nucleus was found to be characteristic of only the longest isoforms, whereas shorter isoforms are present in both the nucleus and the cytoplasm [4]. More recent studies have shown that TCF4 in neurons [17] and colorectal carcinoma cells, in most cases, has a cytoplasmic localization and is correlated with improved survival of colorectal cancer patients [18], suggesting that the protein may have additional, nongenomic functionality. TCF4 mRNAs are ubiquitously expressed in humans, but the levels of TCF4 isoforms vary across different tissues. Low molecular weight isoforms are predominant in the brain: frontal cortex, cerebellum and hippocampus, whereas higher molecular weight isoforms are predominant in the testis and lung [4].

The bHLH domain contains a highly conserved basic region composed of positively charged amino acid residues, allowing it to interact with negatively charged DNA, and the HLH structure responsible for the formation of homo- and heterodimers with other bHLH TFs [19–21]. Upon dimerization, the bHLH domain forms a four-helix bundle with a hydrophobic core, where each monomer contacts an E-box DNA element (5’–CANNTG–3’) half site [20–24]. The C-terminal bHLH domain of TCF4 is the only structurally characterized fragment of TCF4 [24,25]. It has been shown to be involved in the binding of the E-box [20,21,26] in the promoters and enhancers of TCF4-responsive genes, such as the μE5 heavy and κE2 light chain immunoglobulin enhancers [27], an enhancer in the murine leukemia virus SL3-3 genome [16], the rat tyrosine hydroxylase enhancer [28], and the human somatostatin receptor-2 promoter [5]. Mutations associated with a genetic disorder known as Pitt-Hopkins syndrome (PTHS), which is characterized by intellectual disability, distinct facial features, developmental delay, and autonomic dysfunction, mostly occur within the bHLH domain of TCF4 [6,7,29–32]. Interestingly, associations of TCF4 mutations occurring outside the bHLH domain with schizophrenia have been reported; common variants in human *Tcf4* were among the first genes to reach significance in genome-wide association studies of schizophrenia [33], and rare coding *Tcf4* variants outside of the bHLH domain have been identified in individual schizophrenia patients via deep sequencing [34,35].

Despite the expansion of knowledge regarding the function of TCF4, the expression patterns of individual isoforms in tissues, and the consequences of mutations, the understanding of the structure and molecular properties of the TCF4 polypeptide chain remains limited. The only structural report available for TCF4 is related exclusively to the bHLH domain [24]. The *in silico* prediction of the full-length TCF4 structure shown in our previous work suggested that the regions outside the bHLH domain in TCF4 might be disordered [36]. In fact, some TFs have been shown to have a dual, partially disordered nature [37]. The existence of these so-called intrinsically disordered regions (IDRs) is due to the specific amino acid composition, i.e., such regions are enriched in charged residues and depleted in hydrophobic ones. The disordered conformation confers many functional advantages. It is flexible and can quickly switch between different folds or remain unfolded. Disordered regions have also been shown to be susceptible to post-translational modifications. The activation domains of TFs are often disordered [38,39], which significantly increases the range of possible interactions and explains the regulation of protein function.

TCF4 exists in multiple isoforms containing different numbers of activation and repression domains, indicating that the protein is predisposed to interact with a variety of different partners [4,40,41]. TCF4 has been proposed as an important transcription-regulating hub protein [42]. The object of our study was isoform I^-^ of human TCF4. The isoform is expressed predominantly in the brain, according to a publication by Sepp et al. [4]. The lack of fundamental knowledge about the structure of TCF4, prompted us to characterize the full length of the protein, including the region predicted as the IDR. Given that the pivotal function of TCF4 as a transcription factor is binding of DNA through the bHLH domain, we sought to determine whether the molecular properties of this hub protein can be altered upon E-box binding. Several elegant demonstrations of the effects of bHLH binding to specific DNA sequences have been published [43–45]. However, these studies focused only on the protein fragments involved in DNA binding, and systematic studies of full-length bHLH TFs are limited. In this study, we not only are the first to demonstrate the molecular characterization of full-length TCF4 but also present results that highlight the dual nature of the TCF4 structure. The ordered bHLH domain binds the E-box sequence and anchors the protein to a specific genomic location. The regions outside bHLH, which account for 90% of the sequence, are characterized by a state of complete disorder. This, in turn, results in an extended, flexible conformation. Consequently, TCF4 is capable of interacting with multiple partners, thereby enabling the protein to function as a hub transcription factor. Importantly, our observations revealed that DNA interaction does not affect the conformation of the fragments outside the binding region, but E-box binding alters the stability of the TCF4 molecule.

## 4. Materials and methods

### 4.1. *In silico* analysis of the TCF4 primary sequence

In the present study, all the experiments were performed with the human TCF4 isoform I^-^ (UniProt ID P15884-16). It will be referred to as TCF4 in later sections of this article.

Analysis of the TCF4 sequence was performed using bioinformatics tools with default settings. The ProtParam tool (https://web.expasy.org/protparam/) allows the computation of various physical and chemical parameters (e.g., molecular weight, theoretical pI, extinction coefficient). Disorder prediction was performed via the PONDR server (http://www.pondr.com) [46], IUpred3 (https://iupred.elte.hu/) [47–49] and NetSurfP-2.0 (https://services.healthtech.dtu.dk/service.php?NetSurfP-2.0) [50]. Protein backbone dynamics was calculated using DynaMine (http://dynamine.ibsquare.be) [51,52]. Charge-hydropathy analysis was made using the PONDR server [46]. Secondary structure prediction was made by PSIPRED [53]. Example full atom conformations of the TCF4 dimer were generated by ColabFold [54] on the basis of the MMseqs2 homology search and AlphaFold 2.0 [55]. Protein structures were drawn via Discovery Studio 2024 (Dassault Systemes BIOVIA).

### 4.2. Preparation of the expression construct

The cDNA of TCF4 was synthesized *de novo* in GeneArt and optimized via a Gene Optimizer (Thermo Fisher Scientific) to obtain the highest expression level of the synthetic gene in *Escherichia coli*. The optimized sequence was used as a template for PCR, and the primers used for amplification were as follows: TCF4_F: GCGAACAGATTGGTGGCATGatgcaggatggtcatcatagc, TCF4_R: GTGCTCGAGTGCGGCCGC<u>CTA</u>catctgacccatatgattgctt. The obtained insert was subsequently inserted into the pET-SUMO vector (EMBL, Germany) [56] via restriction-free (RF) cloning [57,58]. The sequence of the obtained expression plasmid was confirmed by sequencing.

### 4.3. Protein overexpression for *in vitro* analyses

*E. coli* ArcticExpress competent cells (Agilent Technologies) were transformed with pET-SUMO/TCF4 recombinant plasmids by heat shock and grown overnight at 37 °C on Luria Broth (LB) agar plates supplemented with kanamycin (50 μg/mL) and gentamicin (100 μg/mL). Single colonies were picked and grown in LB media (Invitrogen) supplemented with kanamycin (50 μg/mL) and gentamicin (100 μg/mL) at 37 °C overnight. The main culture in Terrific Broth (TB) media (Invitrogen) supplemented with 50 μg/mL kanamycin was inoculated to a final OD_600_ of 0.1. The culture was incubated at 37 °C and 182 rpm until the OD_600_ reached 0.6, after which the culture was cooled to 16 °C, and protein expression was induced with 0.25 mM isopropyl β-D-1-thiogalactopyranoside (IPTG). The expression continued for 18 h. The bacteria were harvested by centrifugation (5000 × g at 4 °C for 10 min), resuspended in buffer L (20 mM Tris-HCl, 300 mM NaCl, 20 mM imidazole, 5% (v/v) glycerol, 2 mM TCEP, pH 7.5) and stored at −80 °C until purification.

### 4.4. Purification of TCF4

The frozen cell suspension was slowly thawed on ice and supplemented with 20 μg/mL DNase I (Merck), 20 μg/mL RNase A (Sigma Aldrich), 1 mM β-mercaptoethanol (Roth), 200 μg/ml phenyl methyl sulfonyl fluoride (PMSF) (Sigma Aldrich) and 50 μg/mL lysozyme (Sigma Aldrich). The cells were lysed via sonication. The lysates were then centrifuged at 18 000 × g at 4 °C for 1.5 h. The soluble fractions were purified via immobilized metal affinity chromatography (IMAC). The cell lysate was incubated for 1 h at 4 °C with Ni-NTA His•Bind® Resin (Novagen), which had been previously equilibrated with buffer L. Afterwards, the resin was washed with 10 column volumes (cv) of buffer L to remove unbound proteins, then washed with 5 cv of buffer L supplemented with 6 M urea and re-equilibrated with 10 cv of urea-free buffer L. HisTag-SUMO was removed from the fusion protein by cleavage on resin at 4 °C overnight, using 5 cv of buffer L containing 20 μg/mL SUMO hydrolase dtUD1 (EMBL, Germany) [56]. Contrary to expectations, TCF4 was not present in the flow-through fraction after cleavage of HisTag-SUMO. Instead, the protein was released from the resin using 5 cv of buffer E containing imidazole (20 mM Tris-HCl, 300 mM NaCl, 500 mM imidazole, 5% (v/v) glycerol, 2 mM TCEP, pH 7.5). The eluted fractions were pooled, concentrated to a total volume of 1 mL using the Amicon Ultracel-4 centrifugal filter units (Merck Millipore) with a cut-off limit of 30 kDa and injected into the Superdex 200 Increase 10/300 GL column (GE Healthcare Life Sciences) equilibrated with buffer S (20 mM Tris-HCl, 150 mM NaCl, 2 mM TCEP, pH 7.5). The column was operated at room temperature with a flow rate of 0.5 mL/min on an Äkta Avant system (GE Healthcare Life Sciences). Purified TCF4 samples were stored at −80 °C in buffer S. The purity of the protein was estimated densitometrically using ImageJ [59] and was approximately 95% (Additional file 1: Fig. S1A). The molecular mass of recombinant TCF4 was determined via ESI MS at a Mass Spectrometry Laboratory (IBB PAS, Warsaw) (Additional file 1: Fig. S1B).

### 4.5. TCF4 labeling

TCF4 was labeled with Alexa Fluor 488 NHS Ester (Invitrogen). The protein in buffer P (10 mM phosphate, 150 mM NaCl, 2 mM TCEP) was concentrated to 50 μM and incubated with the label at a 1:20 ratio. The conjugation reaction was performed for 1 h at RT, and the pH was adjusted to 8.0. The unbound label was separated from the sample using a Superdex 200 Increase 10/300 column (GE Healthcare). This step also provided exchange of buffers to S. The calculated degree of labeling (DOL) was 1.2 on the basis of the following equation:

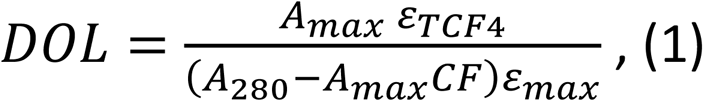

where A_max_ is the absorbance at λ_max_, ε_TCF4_ is the molar extinction coefficient (in M^-1^ cm^-1^) of the pure protein at 280 nm, ε_max_ is the molar extinction coefficient (in M^-1^ cm^-1^) of the dye at its absorbance maximum, and CF is the correction factor.

### 4.6. SDS-PAGE

Protein samples were analyzed by SDS-PAGE using 4%/10% polyacrylamide gels developed in a Tris/Glicyne system [60]. Electrophoresis was performed at a constant current of 20 mA/1 mm gel. The Unstained Protein Molecular Weight Marker (Thermo Fisher Scientific) was used. After electrophoresis, the gels were stained with Coomassie Brilliant Blue R-250 [61].

### 4.7. dsDNA binding assays

To determine the dissociation constant (K_D_), we assumed that double-stranded DNA is bound by the dimeric TCF4 (dTCF4) at a 1:1 stoichiometry on the basis of the known properties of the E-box-binding bHLH transcription factors [62]. The dTCF4:E-box K_D_ was determined by fluorescence polarization (FP), electrophoretic mobility shift assay (EMSA), and fluorescence correlation spectroscopy (FCS) based on the fluorescence of the 6-carboxy-fluorescein (FAM)-labeled double-stranded DNA (dsDNA) probe (FAM-CCGGT<u>CACGTG</u>TCCTA[24], where the E-box canonical sequence is underlined).

#### Fluorescence polarization

FP experiments were performed using a CLARIOstar Plus plate reader (BMG LABTECH) at RT. All measurements were performed in triplicate, using default settings and top optics. Prior to measurements, the dsDNA probe (40 nM) was incubated with increasing amounts of the dTCF4 protein (0–1.5 μM, calculated for the dimer) for 30 min in buffer S with the addition of BSA (final concentration of 0.5 mg/ml) to avoid nonspecific interactions.

#### Fluorescence correlation spectroscopy

Prior to the FCS experiments, the TCF4 protein solution was centrifuged to remove nonspecific aggregates. A FAM-labeled dsDNA probe at 100 nM was incubated with increasing concentrations of dTCF4 (0–2.5 µM per dimer) for at least 30 min. The FCS experiments were performed on a Zeiss LSM 780 with ConfoCor 3 in buffer S in droplets of 25 µl. A single measurement lasted 3 s and was repeated 50 times in series. The series were repeated in 3 independent droplets. The temperature inside the droplet (23 ± 1 °C) was checked after the FCS measurements using a certified calibrated microthermocouple. The structural parameter (*s*) was determined with the AF 488 probe in pure water (D_AF 488_ = 435 μm^2^s^−1^) [63] individually for each microscope slide previously passivated with BSA. The actual solution viscosity of the buffer and solutions at increasing protein concentrations was determined by comparison of the diffusion times for AF 488 in pure water and in the solution at the same instrument calibration. The experiments were performed at an excitation wavelength of 488 nm with a relative argon multiline laser power of 3%, MBS 488 nm, BP 495–555 nm and a damping factor of 10%. The resulting diffusion time (Τ_app_) of the dsDNA probe measured by FCS was the weighted average of the freely diffusing probe and the dTCF4-bound form. The raw measurements were manually curated to eliminate possible nonspecific oligomers or aggregates in the confocal volume. A one-component model of 3D diffusion was used, taking into account the triplet state lifetime of the probe. The FCS data were analyzed by global fitting via Zen2010 software (Zeiss) essentially as described previously [64], and the Τ_app_ values were subsequently corrected for increasing solution viscosity. The total experimental uncertainty of the FCS results was determined according to the propagation rules for small errors [65], taking into account the statistical dispersion of the FCS results and the uncertainty of other experimental values used for calculation of the results.

#### Electrophoretic Mobility Shift Assay

EMSA was performed on the same set of samples used in the FP assays. An aliquot of 10 μl of the reaction mixed with glycerol (final concentration of 20%) and bromophenol blue (final concentration of 1%) was loaded onto 5% native polyacrylamide gels and run at 150 V for 150 min in 0.5 × TBE buffer at 4 °C. Images were captured using the ChemiDoc MP Imaging System (Bio-Rad). DNA probe loss was estimated densitometrically from the captured image (Additional file 1: Fig. S2A) using ImageJ [59].

#### Numerical data analysis of dsDNA binding assays

OriginPro software (version 9.0) and GraphPad Prism (version 6.07) were used to calculate the mean values and standard deviations from the measurements and to perform nonlinear, least-squares regressions. Millipolarization (mP) from the FP experiments and the apparent diffusion time (τ_app_) from the FCS experiments (denoted Y) are described by the following equation:

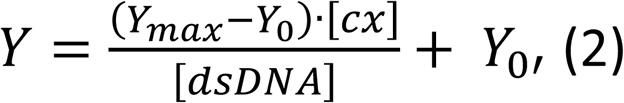

where Y_max_ and Y_0_ are the signals (mP or τ_app_) of the probe in the complex with dTFC4 and in the free state, respectively, and [cx] is the equilibrium concentration of the dTCF4:E-box complex, expressed as a function of the total dTCF4 concentration, according to the following equation:

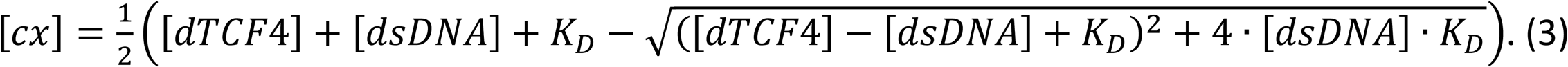

The theoretical curve (eq. 1 + 2) with the free parameters K_D_, Y_max_ and Y_0_ was fitted to the experimental data points. EMSA data were analyzed according to the modified Hill equation, which could compensate for deviations from ideal conditions [66], as follows:

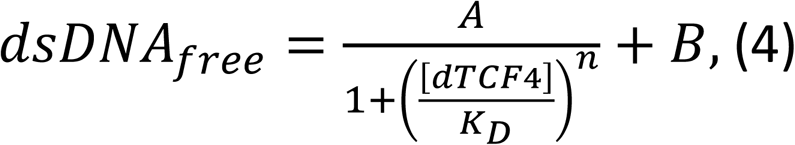

where A and B are coefficients representing the upper and lower plateaus of the titration, respectively, and n is the Hill coefficient. Statistical analysis was performed on the basis of runs tests (P value) and goodness of fit (R^2^), as previously described [67].

We used the K_D_ value determined by the FP technique to calculate the amount of specific dsDNA (CCGGT<u>CACGTG</u>TCCTA, where the E-box canonical sequence is underlined) used in other experiments to compare the properties of free TCF4 and DNA-bound TCF4. In each case, we assumed a molar excess of DNA such that the saturation of dTCF4 with the dsDNA probe was not less than 95%.

### 4.8. Hydrogen/deuterium exchange mass spectrometry

Hydrogen/deuterium exchange mass spectrometry (HDX-MS) was performed on free TCF4 and the TCF4-DNA complex. For measurements of TCF4 in complex with DNA, an E-box was used at twice the molar excess of dsDNA per dTCF4, and samples were incubated on ice for 30 minutes prior to the experiment. Prior to the HDX reactions, nondeuterated fractions of both variants served as sources for peptide lists. For this purpose, the LC-MS analysis was performed with all steps the same as described below for the HDX runs, but in this case, the D_2_O used for exchange was replaced with H_2_O. Peptide identification was performed using ProteinLynx Global Server software (Waters). For the HDX reaction, the starting stock protein concentration was 7 μM. HDX exchange incubations of both variants were performed at five time points: 10 s, 1 min, 5 min, 30 min, and 2.5 h in quadruplicate. 5 μl aliquots of protein stocks were added to 45 μl of deuteration buffer (20 mM Tris DCl, 150 NaCl, 5 mM TCEP, pD 7.45) at room temperature. The final protein concentration in the deuteration reactions was 0.7 μM. The H/D exchange reactions were quenched by transferring the exchange aliquots to precooled tubes (on ice) containing 10 μl of quenching buffer (2 M glycine in 99.99% D_2_O, pD 2.3). After quenching, the samples were immediately frozen in liquid nitrogen and stored at −80 °C until mass spectrometry measurement. The samples were thawed directly prior to measurement and manually injected onto the nano ACQUITY UPLC system equipped with HDX-MS Manager (Waters). Proteins were digested on a 2.1 mm × 20 mm nepenthensin-2 (POROS^TM^) (AffiPro) column for 1.5 min at 20 °C and eluted with 0.07% formic acid in water at a flow rate of 200 μl/min. The digested peptides were passed directly to the ACQUITY BEH C18 VanGuard precolumn, from which they were eluted onto the reversed-phase ACQUITY UPLC BEH C18 column (Waters) using a 10–35% gradient of acetonitrile in 0.01% of formic acid at a flow rate of 90 μl/min at 0.5 °C. The samples were measured on a SYNAPTG2 HDX-MS instrument (Waters) without IMS mode. The instrument parameters for MS detection were as follows: ESI—positive mode; capillary voltage—3 kV; sampling cone voltage—35 V; extraction cone voltage—3 V; source temperature—80 °C; desolvation temperature—175 °C; and desolvation gas flow rate—800 l/h.

Two control experiments were performed to determine the minimum and maximum H/D exchange levels. To obtain the minimum exchange of each peptide (M_min_), 10 µl of a quench buffer was mixed with 45 µl of D_2_O reaction buffer (20 mM Tris DCl, 150 NaCl, 5 mM TCEP, pD 7.45) prior to the addition of 5 µl of protein stock mixture and analyzed by LC-MS. To obtain the maximum H/D exchange level (M_max_), the deuteration reaction was run for 24 h and then quenched with a quench buffer kept on ice. The control experiments were also performed in quadruplicate.

Peptide lists obtained from nondeuterated protein samples were used to analyze the exchange data using DynamX 3.0 software (Waters). The PLGS peptide list was filtered by minimum intensity criteria— 3000 and minimal product per amino acid—0.3. All the raw files were processed and analyzed via DynamX 3.0 software. All MS assignments in Dynamix were inspected manually. The percentage of deuteration [D%] for all peptides was calculated in an Excel file from exported DynamX 3.0 data, based on the Equation 5, which takes into account the minimal and maximal exchange of a given peptide:

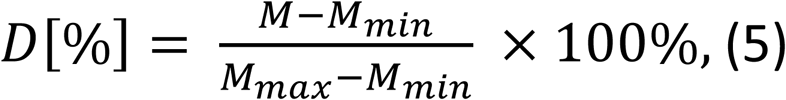

where M is the centroid mass of a given peptide after deuterium uptake, M_min_ is the centroid mass of a peptide with minimal exchange and M_max_ is the centroid mass of a peptide with a maximal exchange. The error bars for the fraction exchanged represent the SDs calculated from at least three independent experiments. The difference in the fraction exchanged (Δ fraction exchanged) was calculated by subtracting the fraction-exchanged values for peptides in the selected state from the values for the same peptides in the control state. Error bars for differences were calculated as the square root of the sum of the variances of the compared states. Student’s t test for two independent samples with unequal variances and unequal sample sizes (also known as Welsh’s t test) was performed to evaluate differences in fraction-exchanged values between the same peptides in two different states. The final data analysis and visualization steps were carried out using the in-house HaDeX software [68].

### 4.9. Native PAGE

Samples of free TCF4 and TCF4 with E-box (the final concentrations of dTCF4 and E-box were 2 μM and 20 μM, respectively) were mixed with glycerol (final concentration of 20%) and bromophenol blue (final concentration of 1%). Then, samples of 20 μl were loaded onto 5% native polyacrylamide gels, and run at 150 V for 40 min in 0.5 × TBE buffer at 4 °C. After electrophoresis, the gels (Additional file 1: Fig. S2B) were stained with Coomassie Brilliant Blue R-250 [61] and analyzed using Image Lab Software (Bio-Rad).

### 4.10. Analytical size exclusion chromatography

The analytical size exclusion chromatography (SEC) experiment was performed on an AKTA Avant FPLC system (GE Healthcare) with a Superdex 200 Increase 10/300 column (GE Healthcare). The column in buffer S was calibrated with a Gel Filtration Marker Kit (MW range 29–700 kDa) (Sigma-Aldrich) according to the manufacturer’s protocol. A total of 75, 50, and 25 μg of TCF4 samples in 100 μl of S buffer were spun down for 10 min at 18000 × g, injected onto the column at a flow rate of 0.5 mL/min and detected by absorbance measurements at wavelengths of 280 and 260 nm. An analogous set of samples at the same concentrations in the presence of E-box dsDNA (twice the molar excess of dsDNA to dTCF4) was then spun down for 10 min at 18000 × g, injected onto the column at a flow rate of 0.5 mL/min and detected by absorbance measurements at 280 and 260 nm wavelengths. E-box dsDNA was also injected onto the column at a concentration of 10 μM. All the injections were performed in triplicate.

### 4.11. Analytical ultracentrifugation

Sedimentation-velocity analytical ultracentrifugation (SV-AUC) experiments were performed on a Beckman Coulter Proteome Lab XL-I ultracentrifuge (Beckman Coulter Inc.) using AN-60Ti 4-hole rotor, a 12 mm path length, a charcoal-filled double sector Epon® centerpiece and quartz windows. The scans were collected at 20 °C and a rotor speed of 50 000 rpm. TCF4 was analyzed at 0.25, 0.50 and 0.75 mg/mL (2.5, 5.0 and 7.5 μM dTCF4 respectively) without and with dsDNA at 5.0, 10.0 and 20.0 μM, respectively, in buffer S with a detection wavelength of 280 nm. The density (1.005 g/mL) and viscosity (0.010213 mPa × s) were calculated using SEDNTERP [69]. The partial specific volume (vbar) of the protein (0.7131 mL/g) was determined from the sequence using SEDNTERP [70], and the ̅ value of the dTCF4:dsDNA complex (0.697 mL/g) was calculated as the weighted average for the protein and DNA [71], where the weights were the MWs of dTCF4 and dsDNA. The time-corrected data were analyzed with SEDFIT software (version 16.1) using the built-in continuous sedimentation coefficient distribution model c(s). The maximum entropy regularization of the models was set to a confidence level of 0.68 [72,73].

Analyses were performed using three built-in models: the continuous sedimentation coefficient distribution c(s) with a monomodal frictional ratio (f/f_0_) value, the continuous c(s) with a bimodal f/f_0_ distribution, and the continuous c(f/f_0_) distribution. In the continuous c(f/f_0_) model, the method for determining the f/f_0_ and hydrodynamic radius (R_h_) values and their errors for an IDP assumes a known molecular weight (MW). Therefore, the MWs were set to known values of 96.5 kDa and 106.4 kDa for dTCF4 and the dTCF4:dsDNA complex, respectively. First, the s distribution was calculated for different f/f_0_ values.

For the s range corresponding to the dimer size (R_h_) and shape (f/f_0_), the distribution was plotted against the MW (Additional file 1: Fig. S3). The f/f_0_ value, which is the best estimate of the dimer shape, was determined using the f/f_0_ value, for which the vertical line indicating MW of the dimer intersected the f/f_0_ distribution in the middle. The uncertainty of the f/f_0_ value (Δf/f_0_) is the difference between the best f/f_0_ value and the nearest analyzed f/f_0_ value. Next, R_h_ was plotted as a function of MW for the best f/f_0_ and two closest f/f_0_ values. R_h_ is the value that intersects the R_h_(MW) function for the best f/f_0_, and ΔR_h_ is the difference between R_h_ and the value determined for the neighboring f/f_0_.

### 4.12. Circular dichroism spectroscopy

Circular dichroism (CD) spectra were recorded using a Jasco-815 spectropolarimeter (Jasco Inc.) equipped with a Peltier temperature controller (CDF-426S/15). The spectra were collected in the spectral range of 195–300 nm at a scanning speed of 50 nm/min at 20 °C, D.I.T. of 2 s and a 1 nm bandwidth. The data pitch was 0.5 nm. The spectra were measured in 5 accumulations using 1 mm path length quartz cuvettes, and the concentration of dTCF4 was 0.5 mg/mL (5 µM). An E-box was applied at twice the molar excess of dsDNA over dTCF4, and the samples were incubated on ice for 30 minutes before measurement. All the spectra were corrected for the effect of buffer and converted to molar residual ellipticity units [74]. The mean residue weight (MRW) for TCF4 I^-^ is 106.65 Da. The secondary structure content was estimated using BeStSel deconvolution software [75–79].

Temperature-dependent denaturation was monitored by following the changes in ellipticity in the range of 195–260 nm by increasing the temperature from 20 to 95 °C and then decreasing it from 95 to 20 °C at a constant rate of 1 °C/min. The melting temperature was determined by fitting a sigmoidal (Boltzmann) curve to the points of the ϴ_MRW_-temperature relationship for a wavelength of 220 nm. The fitting was performed using OriginPro software (version 9.0).

### 4.13. Thermal shift assays

Thermal shift assays were performed using QuantStudio™ 5 Real-Time PCR System (Thermo Fisher Scientific). Each sample contained 5 × SYPRO® Orange Protein Gel Stain (Merck). Stock concentrations of protein, SYPRO® probe and E-box DNA were prepared in buffer S. The assays were performed for the samples containing dTCF4 (5 μM) with and without dsDNA, dsDNA alone, and SYPRO® Orange as a control sample. An E-box was applied at a 2-fold molar excess of dsDNA over dTCF4. All the samples were prepared in 3 replicates. The measurements were performed in a 96-well plate. The samples were incubated on ice for 30 minutes before measurement. The temperature was subsequently increased from 25 to 99 °C at a rate of 0.05 °C/s. The optical filter used was 470 ± 15 nm for excitation and 586 ± 10 for emission. The curves were then averaged for each species and corrected for the temperature exponential decay of the fluorescence intensity on the basis of the correction curve determined from the temperature fluorescence quenching of pure SYPRO® Orange. The melting point was then calculated from the first derivative of the corrected temperature-dependent fluorescence curve as the maximum absolute value of this derivative.

## 5. Results

### 5.1. *In silico* analyses suggest that TCF4 is mostly disordered

Our previous *in silico* predictions of the TCF4 structure suggested that regions outside the bHLH domain in TCF4 may be disordered. Here, we present the results of a detailed analysis to determine the propensity of TCF4 (I^-^) to adopt an intrinsically disordered structure. IDPs are characterized by a low overall hydrophobicity and a large net charge, so it is possible to categorize them using a charge-hydropathy plot [80]. The charge-hydropathy plot prepared for TCF4 (Fig. 1A) shows that this protein is located slightly below the boundary and is grouped with the ordered set of proteins. This may be due to the presence of a locally ordered structure — the well-studied, highly conserved bHLH domain. However, it should be noted that using only charge-hydropathy plot does not give certain results, and as presented in Fig. 1A there are many known examples of proteins documented as disordered whose location also suggests their ordered nature.

**Fig. 1.**
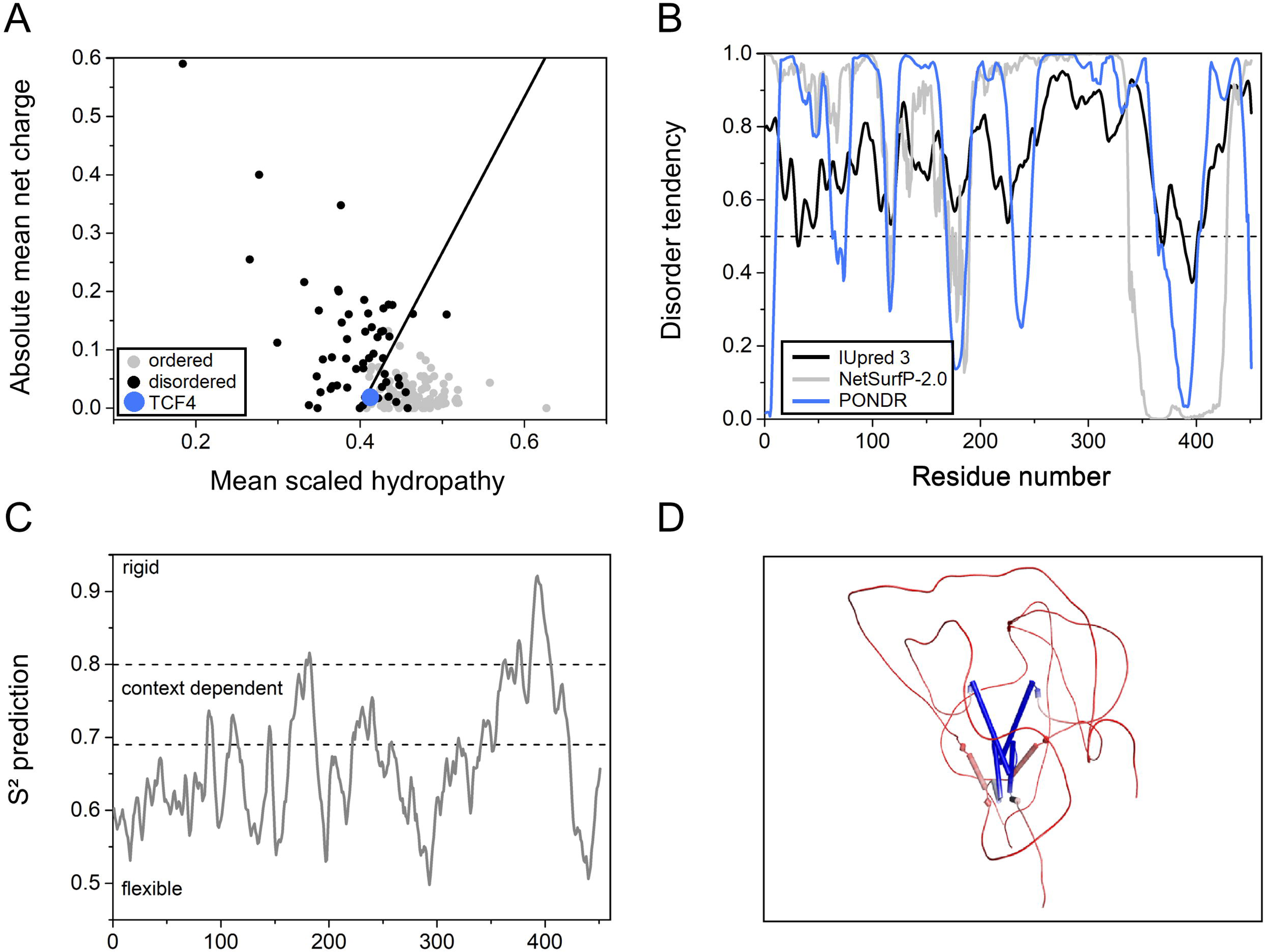
*In silico* analyses of TCF4. (A) Charge-hydropathy plot of fully ordered proteins (gray dots), fully disordered proteins (black dots) and TCF4 (blue dot). The black line represents the boundary between the ordered and disordered proteins. (B) Prediction of the degree of the disorder in the TCF4 sequence calculated from the primary structure. The results using 3 independent predictors are shown: IUpred 3 (black line), NetSurfP-2.0 (gray line), and PONDR (blue line). A score above 0.5 indicates a high probability of disorder. (C) Prediction of protein backbone dynamics using DynaMine. An S^2^ value higher than 0.8 indicates high rigidity of the protein backbone, whereas an S^2^ value lower than 0.69 indicates high flexibility, which is typical for disordered segments. Values between 0.69 and 0.8 are characteristic of the context-dependent structural organization of polypeptide chains. (D) Example TCF4 dimer conformation generated via ColabFold [54] based on the MMseqs2 homology search and AlphaFold 2.0 [55] for visualization purposes [81]. A schematic protein representation colored according to the average isotropic displacement, from low (blue) to high values (red); α-helices are shown as cylinders.

To investigate to what extent TCF4 could reveal features of IDPs, we used independent predictors of disorder: PONDR, IUpred3, PSIPRED, and NetSurfP-2.0. The results obtained with all the algorithms are consistent and indicate the presence of extensive disorderness, of about 80%. The remainder of the sequence consists mainly of a fragment containing the bHLH domain (residues 348–401) (Fig. 1B), and according to the NetSurfP-2.0 prediction results, it is involved in the formation of helices (Additional file 1: Fig. S4A). To predict the dynamics of the protein backbone, we used DynaMine, which defines the presence of flexible regions [51,52]. The more flexible the region is, the greater the probability of the existence of the disordered structures [51,52]. The TCF4 sequence was mostly identified as flexible/disordered, as indicated by values less than 0.69. Context-dependent (values between 0.69 and 0.8) and rigid/ordered regions (values above 0.8) mainly cover the bHLH domain (Fig. 1C). Considering the crystal structures of the TCF4 bHLH domain obtained for both the DNA-free and DNA-bound forms of TCF4 [24], we assumed that the protein exists as a dimer and used a dimeric sequence to simulate its conformation. The results obtained via ColabFold [54] based on the MMseqs2 homology search and AlphaFold 2.0 [55] (Fig. 1D) suggest the possibility of the existence of an additional α-helical fragment in the region of 167–183 of the long N-terminal IDR, which is consistent with the PSIPRED and NetSurfP-2.0 predictions of the helical propensity for the 166–187 fragment (Additional file 1: Fig. S4B) as well as with a decreased disorder tendency (Fig. 1B) and increased rigidity (Fig. 1C) for this sequence region.

The results of the *in silico* analyses suggested that TCF4 might be mainly disordered, but as mentioned earlier, the experimental results do not always match the predictions. For this reason, we decided to conduct *in vitro* experiments to analyze the molecular properties of TCF4. Since TCF4 is a DNA-binding protein, we decided to perform experiments on both DNA-free TCF4 and TCF4 in complex with a specific E-box sequence.

### 5.2. dTCF4 binds the canonical E-box sequence *in vitro*

As indicated above, in this study we focused the differences between TCF4 structures in the free state and in the DNA-bound state. Therefore, it was especially important to obtain a pure TCF4 preparation that did not contain DNA from bacterial cells. During the purification process, we found that to achieve this goal, it was necessary to use a buffer containing 6 M urea for washing the IMAC column with the bound 6xHis-SUMO-TCF4 fusion protein. By omitting this step, the protein preparation was significantly contaminated with bacterial nucleic acids (Additional file 1: Fig. S5).

To test the DNA-binding ability of TCF4, we used a double-stranded fluorescently labeled DNA probe that contained the E-box canonical sequence. We determined the dissociation constant (K_D_) via a fluorescence polarization (FP) assay, electrophoretic mobility shift assay (EMSA) and fluorescence correlation spectroscopy (FCS). To determine the K_D_, we assumed that dsDNA is bound by the dimeric protein (dTCF4) at a 1:1 stoichiometry on the basis of the known properties of the E-box-binding bHLH transcription factors [62].

The results of the FP binding assay are shown in Fig. 2A. The K_D_ value obtained using the FP-based method was determined to be 0.12 ± 0.03 μM. The same set of samples was used to perform EMSA (Fig. 2B). After the EMSA results were acquired, the relationship between the amount of unbound fluorescent probe and the concentration of dTCF4 was determined densitometrically. Numerical analysis of the relative integrated intensity of the bands revealed that the K_D_ value obtained via EMSA was 0.327 ± 0.014 μM.

**Fig. 2.**
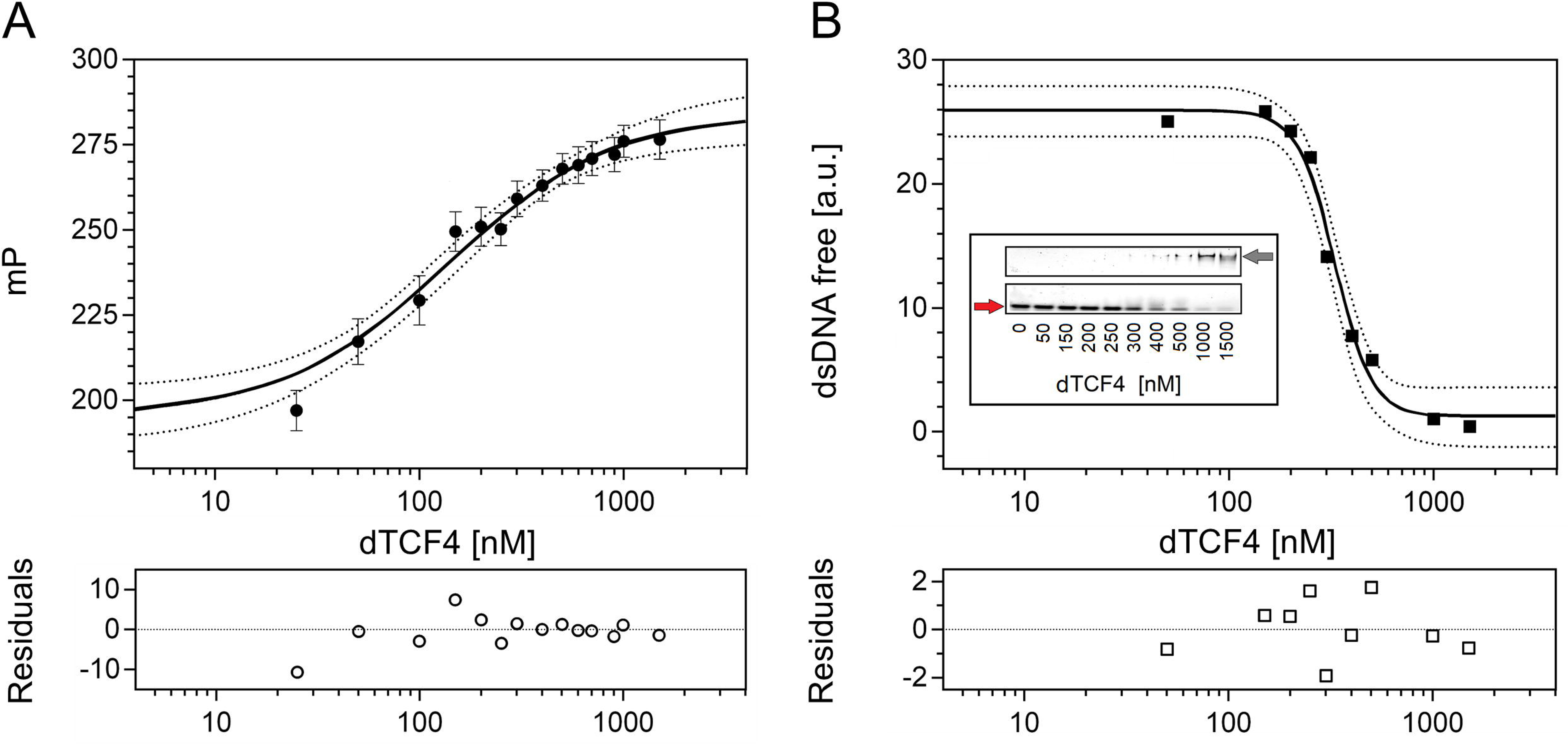
Determination of the dTCF4:E-box K_D_ using the FP assay and EMSA. (A) Data from the FP assay. The black circles are the average millipolarization (mP) values from three measurements, shown with the standard deviation; the binding curve was fitted to the experimental points according to Eq. 2 and Eq. 3 (see the Materials and methods section). (B) Loss of the free fluorescent dsDNA probe determined from the intensity of the bands on the polyacrylamide gel after EMSA (inset); the red and gray arrows show the free and dTCF-bound dsDNA probes, respectively; the binding curve was fitted according to Eq. 4. The accuracy of the binding assays was assessed on the basis of the 95% CI (dotted lines) and fitting residuals (bottom panels).

Next, we performed FCS experiments. The autocorrelation curves show a systematic shift toward longer lag times in the presence of higher amounts of dTCF4 (Fig. 3A). On the basis of the FCS curves, we calculated the apparent diffusion times (τ_app_) of the probe at different protein concentrations. The analysis of the increasing τ_app_ value as a function of the dTCF4 concentration yielded K_D_ = 0.33 ± 0.11 μM (Fig. 3B). The diffusion time of the dTCF4:E-box complex determined from FCS was τ_max_ = 390 ± 30 μs, which corresponds to a hydrodynamic radius R_h_ = 68 ± 6 Å.

**Fig. 3.**
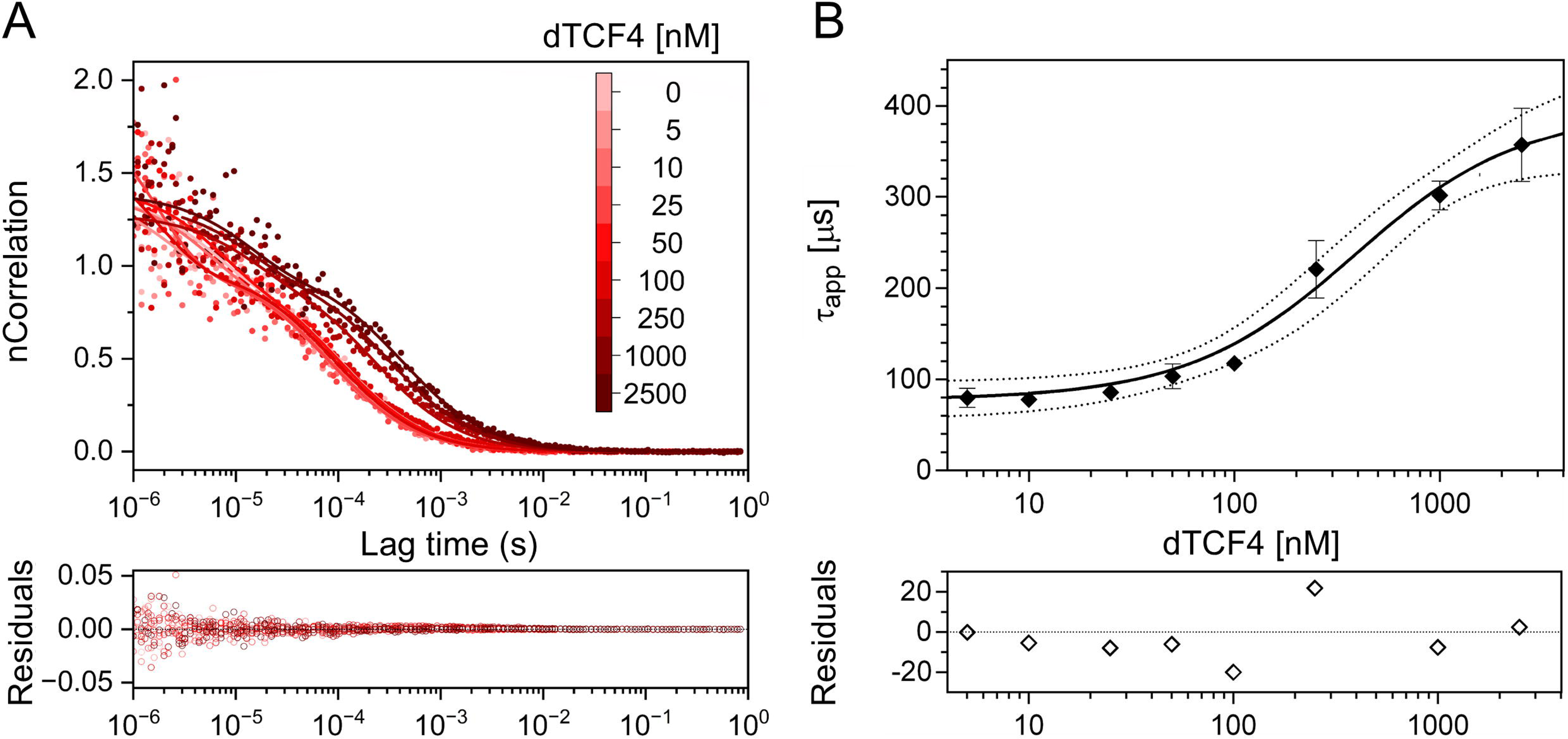
Determination of the dTCF4:E-box K_D_ using FCS. (A) Normalized FCS data (dots) and autocorrelation curves (solid lines) for the FAM-labeled dsDNA probe at 100 nM titrated with increasing concentrations of dTCF4, marked by red intensity (upper panel), together with the corresponding nonnormalized FCS fitting residuals (bottom panel). (B) The binding curve was fitted to the experimental points according to Eq. 2 and Eq. 3 (see the Materials and methods section); the accuracy of the binding assay was assessed on the basis of the 95% CI (dotted lines) and the fitted residuals (bottom panel).

The results of the binding assays for the dTCF4:E-box complex formation are presented in Table 1. All the approaches used showed that the affinity of dTCF4 for the E-box-containing dsDNA probe is submicromolar. The K_D_ value was quantitatively consistent between the EMSA and FCS measurements. The FP results might even indicate a slightly higher binding affinity within the complex but the R^2^ value of the fit is worse because of the limited number of points at lower concentrations. Notably, however, the K_D_ value for the interaction of the E-box with the TCF4 bHLH domain was shown to be sensitive to ionic strength and the presence of a crowding agent [24]. Considering that each of the binding assays was performed under different conditions, we can conclude that the K_D_ values we obtained via the three methods are semiquantitatively concordant, ranging from 0.2–0.3 μM (0.3–0.7 μM, as per monomer).

**Table 1.** Dissociation constants (K_D_) with their 95% confidence intervals (95% CI) and goodness of fit parameters (R^2^ and P value) for dTCF4 binding to a dsDNA probe containing an E-box.

| | $K_D$ (μM)* | $R^2$ | P value |
| --- | --- | --- | --- |
| **FP** | $0.12 \pm 0.03$ | 0.973 | 0.514 |
| **EMSA** | $0.327 \pm 0.014$ | 0.989 | 0.405 |
| **FCS** | $0.33 \pm 0.11$ | 0.983 | 0.643 |

\* $K_D$ calculated vs. TCF4 monomer concentration are $0.26 \pm 0.05$ , $0.65 \pm 0.03$ , and $0.76 \pm 0.23$ μM from FP, EMSA, and FCS, respectively.

We used the K_D_ determined by the FP method to calculate the amount of DNA used in the techniques for comparing free TCF4 with DNA-bound TCF4, which are described further in the article.

### 5.3. TCF4 is almost entirely disordered except for the canonical bHLH domain, which binds the E-box sequence

To directly map the IDRs in TCF4, we utilized hydrogen/deuterium exchange mass spectrometry (HDX-MS). In this technique, the rate of chemical exchange between deuterons in solution and hydrogens on the amide groups of a protein is determined by mass spectrometry. Amide groups deeply buried in the core of a protein or those involved in intramolecular bond formation are characterized by a very low rate of hydrogen exchange. In contrast, amides from the dynamic disordered fragments exchange hydrogens very rapidly. This difference can be used to identify IDRs in a given protein [82,83]. The results of HDX-MS analysis for free TCF4 after 10 s and 150 min are shown in Fig. 4A. The entire sequence, except for the region covering the bHLH domain (residues 360–405) is characterized by efficient, almost total H/D exchange after 10 s of the reaction. After 150 min, the fractional deuterium uptake of peptides containing the bHLH domain increased, indicating the dynamics of this protein fragment.

**Fig. 4.**
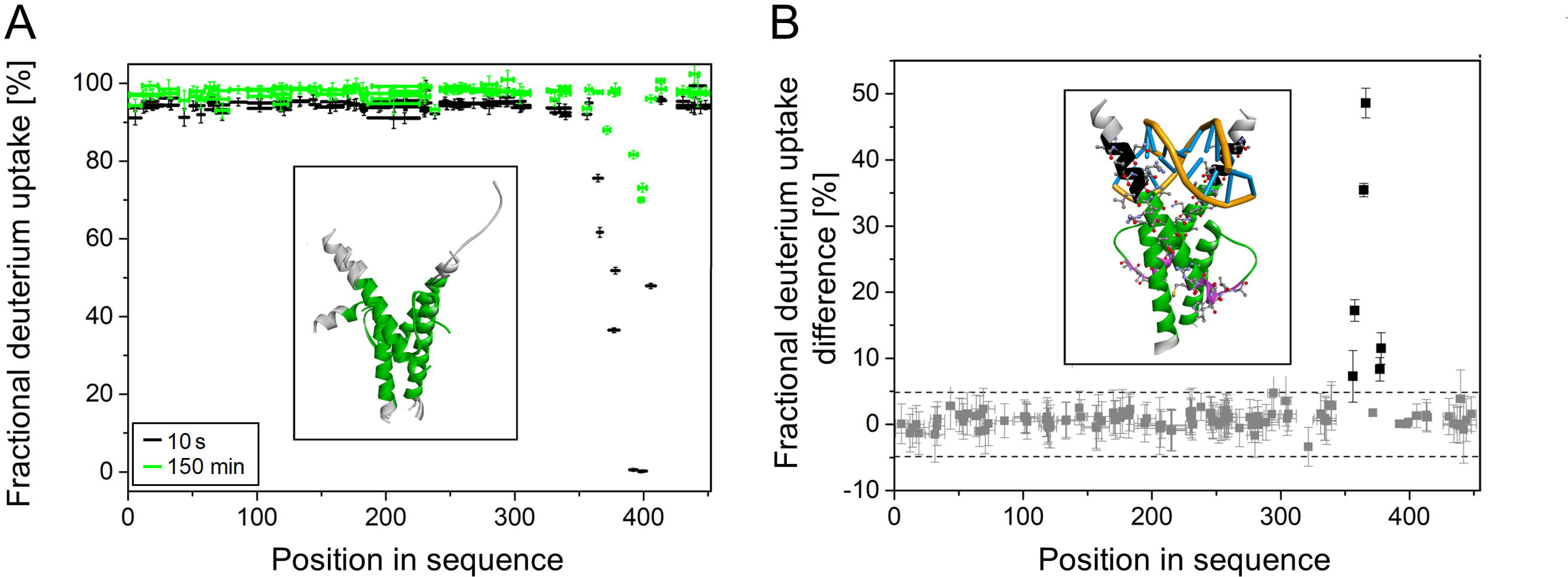
HDX-MS analysis of TCF4. (**A**) Fractional deuterium uptake of free dTCF4 peptides. The horizontal bars correspond to individual peptides generated by the digestion of TCF4 with nepenthensin-2. The vertical bars indicate the measurement error. The black color represents the results of the measurements after 10 seconds, and the green color represents the results after 150 minutes. Inset: Conformations of the *apo* dTCF4 bHLH domain (pdb 6OD3, 6OD4, [24]) with the residues protected from the immediate HDX (360–405) marked green. (B) Fractional deuterium uptake difference at 10 s between the free and dsDNA-bound forms of dTCF4. The gray squares located within the dashed lines correspond to peptides for which the fractional deuterium uptake difference between the two dTCF4 forms is insignificant (within the 98% CI). The black squares above the upper dashed line correspond to peptides of free dTCF4 that are significantly protected upon the binding. Inset: dTCF4 bHLH domain bound to the E-box (pdb 6OD4, [24]) with the residues more protected in the complex (352–367 and 372–380) shown as balls and sticks; the residues protected in the complex but exposed in the *apo* form (352–359) are marked in black; the residues more protected in the complex, but without direct contacts with DNA, are marked in magenta.

The next step was to perform an analogous experiment for TCF4 in complex with the dsDNA probe. Compared with the free form of TCF4, we found that TCF4 bound to dsDNA contained an additional region of slower exchange. These differences were observed for the peptides involving amino acid residues 352–367 (ANNARERLRVRDINEA) and 372–380 (GRMVQLHLK) (Fig. 4B), and their sequences are listed in Table 2. This finding indicates that the residues 352–359 (ANNARERL) preceding the canonical HLH domain are protected from the immediate HDX while the protein is bound to a specific DNA sequence.

**Table 2.** Peptides that differ in fractional deuterium uptake depending on DNA binding by TCF4.

| Sequence | Position |
| --- | --- |
| ANNARER | 352–358 |
| NARERL | 354–359 |
| RVRDINEA | 360–367 |
| DINEA | 363–367 |
| GRMVQLHLK | 372–380 |
| MVQLHLK | 374–380 |

A 98% confidence interval was used for the Student's test.

After binding of the DNA, the results of the HDX-MS experiment revealed a slower exchange in the DNA-binding region of TCF4 (Fig. 4B inset, black residues), which could be expected for the complex on the basis of the crystal structures [24]. In contrast, a surprising result was obtained for the second region of TCF4, which is located at the C-terminus of the first α-helix and at the beginning of the loop of the bHLH domain (residues 372–380). The region is not directly engaged in the interaction with the E-box (Fig. 4B inset, magenta residues), however, it was HDX-protected upon the complex formation. This suggests that the E-box binding leads to increased compactness of the dTCF4 hydrophobic core. Surprisingly, no effect of DNA binding was observed on disordered regions of TCF4, except for the basic region of the bHLH domain.

We also performed UV CD spectroscopy measurements of both TCF4 forms. The recorded spectra differed only slightly, suggesting only a transition of the basic region of the bHLH domain from disordered to ordered toward the α-helical fold after DNA binding (Additional file 1: Fig. S6). The results obtained with CD indicate, in line with the HDX-MS results and crystal structures [24], that there is no significant effect of E-box DNA binding on the overall protein structure.

### 5.4. TCF4 appears exclusively as a dimer regardless of the presence of an E-box

To comprehensively analyze the impact of DNA on TCF4, we decided to investigate the effect of DNA binding on the hydrodynamic properties of TCF4. Our results presented above revealed a significant degree of TCF4 disorder. Given that TCF4 is a relatively large IDP, we were aware that in contrast to globular proteins, MW of TCF4 cannot be calculated from its R_h_. Considering the crystals of the dimeric bHLH domain obtained for both the DNA-free and DNA-bound forms of TCF4 [24], we assumed that full-length TCF4 occurs in dimeric form both in the *apo* and *holo* states. We carried out a series of experiments (native PAGE, SEC, and SV-AUC) to test the hydrodynamic properties of the molecules in their free and DNA-bound states. The first experiment we performed in this series was native electrophoresis, the results of which are shown in Fig. 5, inset. The resulting image after DNA binding differs slightly from that of free TCF4. The band moved faster in the polyacrylamide gel, probably because the TCF4 complex gained an additional negative charge carried by the DNA. In both types of samples, we observed single bands demonstrating single populations of molecules with similar properties under native PAGE conditions.

**Fig. 5.**
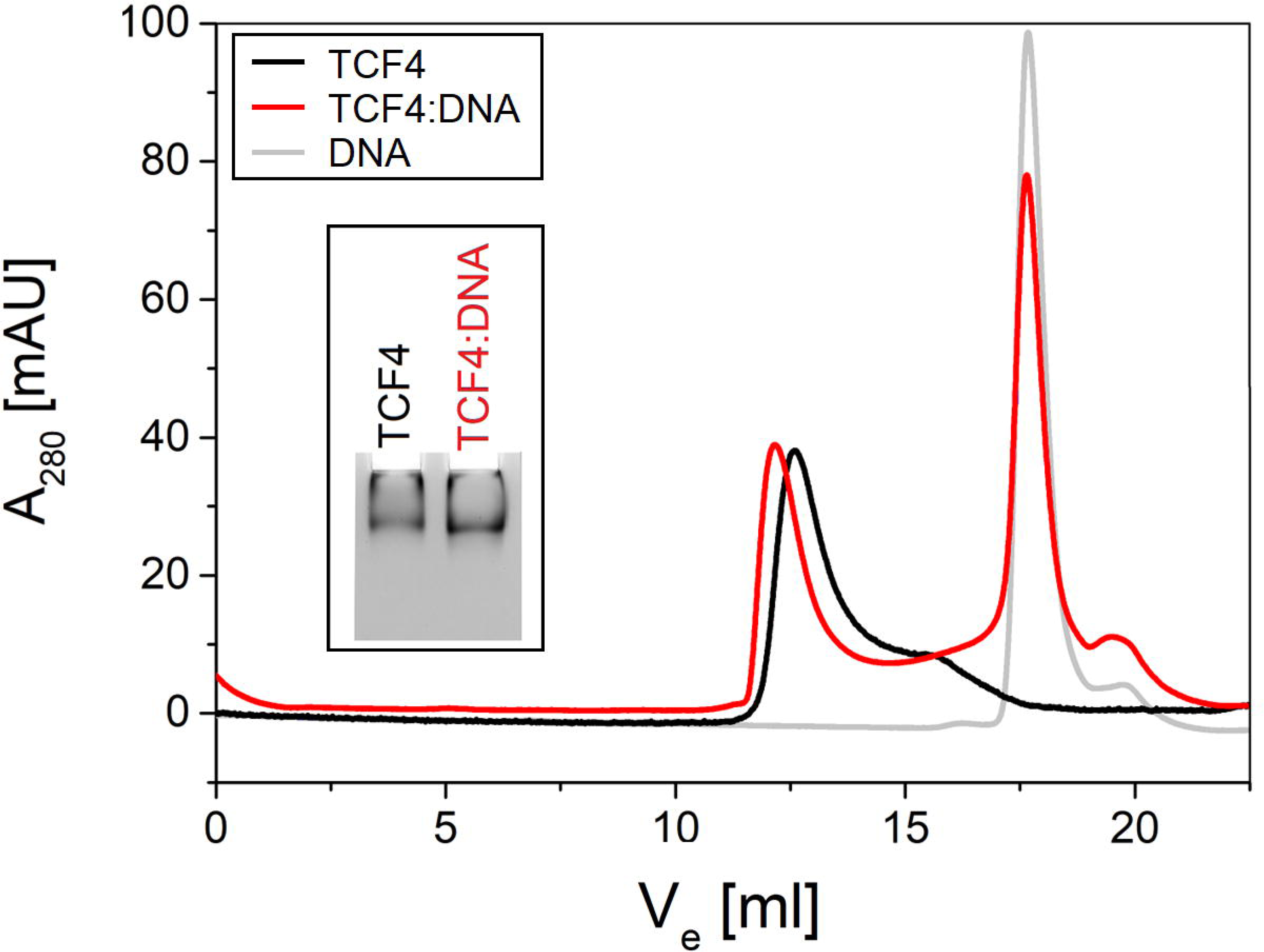
Determination of TCF4 population homogeneity using native PAGE and SEC. Overlaid representative chromatograms at 280 nm from SEC of free TCF4 (black), DNA-bound TCF4 (red), and free DNA (gray). The absorbance values for the free form of TCF4 were multiplied by a factor of 4.5, for simplified readability of the graph. Inset: Native PAGE of free TCF4 and TCF4 bound with DNA performed on 5% polyacrylamide gel.

Next, we performed an analytical SEC of free and DNA-bound TCF4. In this technique molecules are separated on the basis of their hydrodynamic properties. The free form of TCF4 showed a single major peak (Fig. 5, black line). Chromatography of the TCF4 sample with DNA revealed two single peaks, the first with an elution volume approximating the major peak coming from the free form of TCF4 and the second indicating the presence of significantly smaller molecules (Fig. 5, red line). By performing a control free DNA chromatography experiment, we classified the second peak as a signal from unbound DNA (Fig. 5, gray line). TCF4 elution profiles indicate the existence of a population of TCF4 molecules with similar properties in each of the samples analyzed, regardless of the presence of DNA. On the basis of these findings, upon binding, the hydrodynamic properties of the TCF4:DNA complex barely change in relation to those of the *apo* form of TCF4.

To further test the conformational changes in TCF4 induced by DNA binding, we performed an SV-AUC experiment. Representative sedimentation profiles are presented in Fig. 6A, 6B. The protein was analyzed at three different concentrations in the absence or presence of the ligand in the form of E-box DNA. In this analysis, we focused on determining the changes in the overall shape of dimeric TCF4 in relation to its *apo* and *holo* forms. We applied the c(f/f_0_) model and determined the R_h_ values of both forms. To do so, we first determined the sedimentation coefficient (s) distribution for both TCF4 forms with the c(f/f_0_) model (Additional file 1: Fig. S7). The best match of s distribution was obtained for *apo* dTCF4 for f/f_0_ = 2.41 and for dTCF4:dsDNA for f/f_0_ = 2.37 (Fig. 6C, 6D). Our method assumed a known molecular weight (i.e., 96.5 kDa for dTCF4 and 106.4 kDa for the TCF4:DNA complex). The dimer size (R_h_) and shape (f/f_0_) distributions were plotted with respect to the MW (Fig. 7A, 7B). This method allowed us to determine the R_h_ of both forms of dTCF4. The R_h_ of the *apo* form was 72.7 ± 1.1 Å, whereas for dTCF4 in the presence of DNA, it was 67.5 ± 1.1 Å. These results indicate that once the dTCF4 binds to DNA, the complex becomes more compact.

**Fig. 6.**
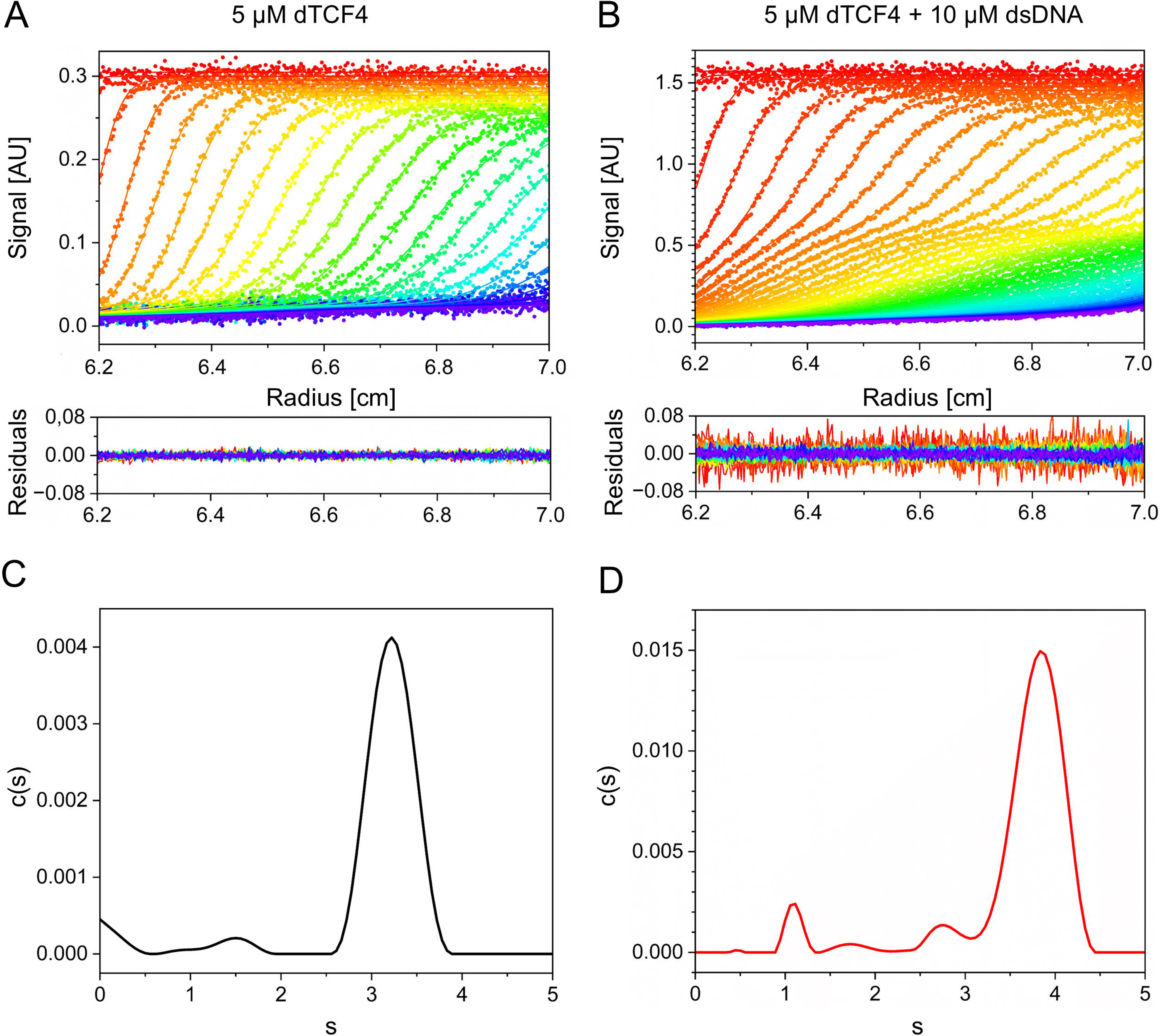
SV-AUC analysis of free and DNA-bound TCF4. Sedimentation profiles of (A) dTCF4 sample with no DNA and (B) with specific DNA. Sedimentation coefficient (s) distributions of (C) free dTCF4 and (D) dTCF4 with specific DNA calculated using the c(f/f_0_) model for f/f_0_ = 2.41 and f/f_0_ = 2.32, respectively. The presented data correspond to dTCF4 analyzed at 5 µM and 5 µM dTCF4 in the presence of 10 µM specific DNA.

**Fig. 7.**
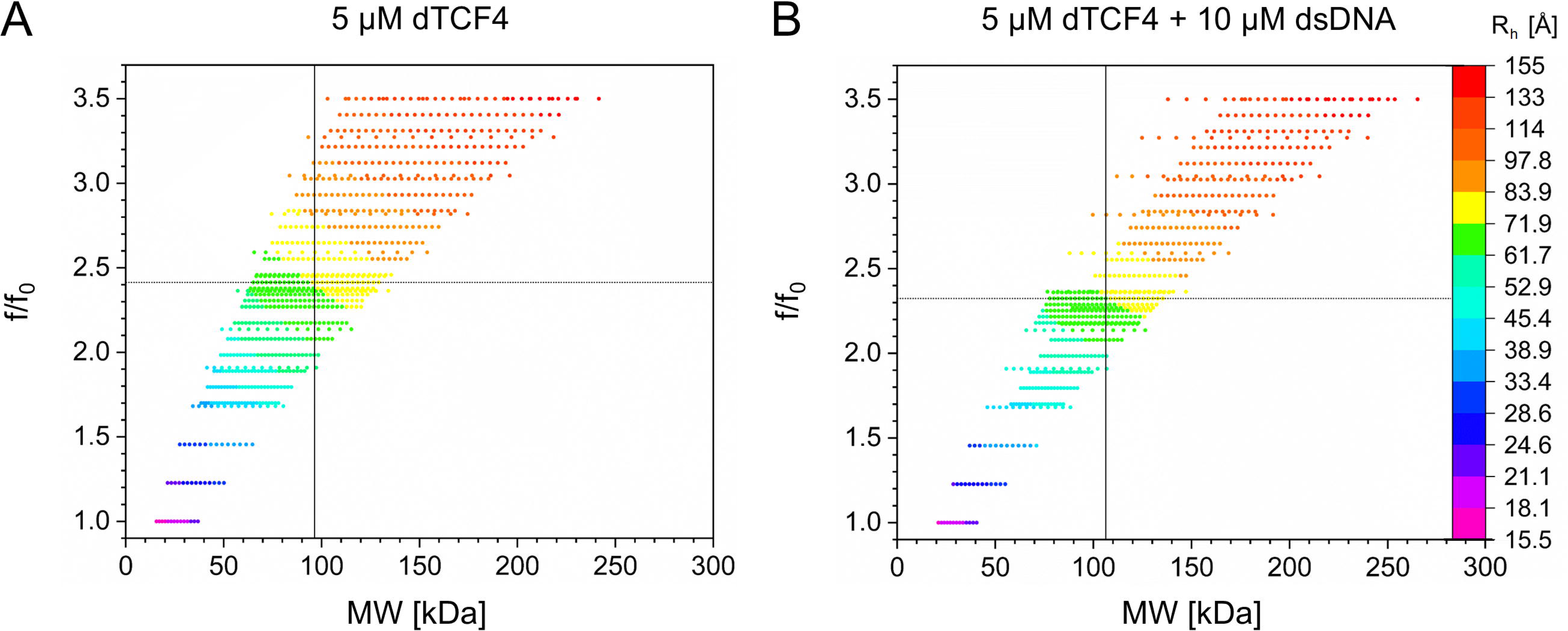
Size (R_h_) and shape (f/f_0_) parameters of TCF4 determined by the SV-AUC. Size (R_h_) and shape (f/f_0_) distributions shown with respect to molecular weight (MW) for (A) 5 µM dTCF4 with no DNA and (B) with 10 µM DNA (= 0.697 mL/g). Solid line, (A) dTCF4 and (B) dTCF4:dsDNA complex MW values; dashed line, f/f_0_ values for the corresponding MWs.

### 5.5. DNA binding influences the stability of the TCF4 molecule

It has been reported that some proteins stabilize their structure after binding ligands [84] and it can correlate with the changes in protein flexibility [85]. To investigate the effect of DNA binding to TCF4 on the thermal stability, free and DNA-bound proteins were again analyzed using a thermal shift assay and CD experiments at different temperatures. In the thermal shift experiment, we observed a prominent peak in the fluorescence maximum for DNA-bound TCF4 in the temperature range studied. For free TCF4, the fluorescence was high from the beginning of the heating process, and then, its value increased only slightly (Fig. 8A). The SYPRO® Orange probe exhibits fluorescence after interacting with hydrophobic residues of the protein. The high fluorescence value from the beginning of the process and the lack of the typical transition for free TCF4 may be caused by the exposed hydrophobic residues in the native form of free TCF4. This means that in the *apo* state, TCF4 has exposed regions of the hydrophobic character. Most likely, not only the basic region preceding the HLH is not folded but also the N-terminal part of the HLH domain. Interestingly, in the DNA-bound state, the increase in the fluorescence values observed with increasing temperature is sharp and rapid. This suggests that after DNA binding, part of TCF4 folds, leading to the formation of a hydrophobic structure that can bind SYPRO® Orange before a heat-induced conformational transition occurs.

**Fig. 8.**
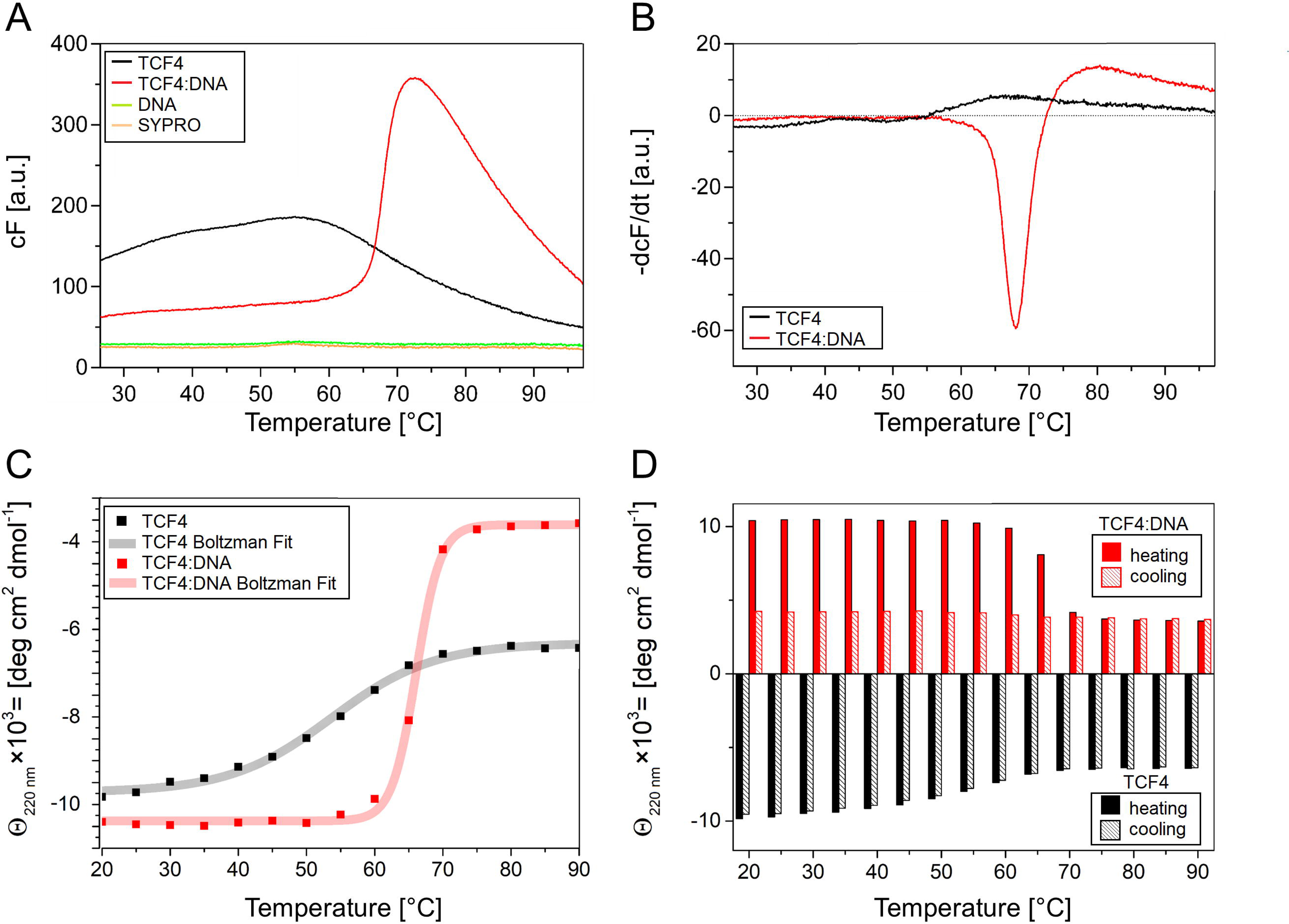
Investigation of the effect of DNA binding on the thermal stability of TCF4. (A) Curves of the dependence of the fluorescence of the SYPRO® Orange probe on the temperature. Black represents the curve for free TCF4, red represents TCF4 in complex with DNA, green represents free DNA, and orange represents free SYPRO® Orange. (B) Curves of the first derivatives of fluorescence as a function of temperature. The black color indicates the result for free TCF4, and the red color indicates TCF4 in complex with DNA. (C) Curves of the dependence of the molar-residual ellipticity at 220 nm on the temperature (heating process). Black - points and the fitted sigmoidal curve for free TCF4, and red - TCF4 in complex with DNA. (D) Bar graph of the dependence of the molar-residual ellipticity at 220 nm on temperature. The black color indicates the values of molar-residual ellipticity for free TCF4 (solid fill— heating process, dashed fill — cooling process), and the red color indicates TCF4 in complex with DNA (solid fill — heating process, dashed fill — cooling process). The ellipticity values for DNA-bound TCF4 are shown as absolute values for simplified readability of the diagram.

The determination of the first derivative of the fluorescence as a function of temperature allowed the determination of the melting point of TCF4 in complex with the DNA. The melting temperature was approximately 68 °C. In the case of free TCF4, the melting point was approximately 50 °C (Fig. 8B).

To explore the influence of DNA binding on the stability of TCF4, we decided to use an alternative technique, CD spectroscopy. We recorded CD spectra in the temperature ranges of 20–90 °C and 90– 20 °C for free TCF4, and for DNA-bound TCF4. We then plotted the dependence of the ellipticity at 220 nm on temperature during the heating process and fitted sigmoidal curves to the resulting series of points (Fig. 8C). The melting temperatures were approximately 53 °C for free TCF4 and 66 °C for DNA-bound TCF4, which is consistent with the results obtained in the thermal shift assay. We also noted that the decrease in the absolute ellipticity value at 220 nm for free TCF4 was much lower than that for DNA-bound TCF4 (Fig. 8C).

This CD spectroscopy analysis identified an interesting case of free TCF4 behavior, which was revealed by monitoring the ellipticity at 220 nm during cooling. We observed that the absolute value of ellipticity at 220 nm, which decreased with heating, completely returned to its initial value during the cooling process. This effect was not observed for DNA-bound TCF4, which exhibited a typical irreversible transition (Fig. 8D).

## 6. Discussion

In this study, we present the results of a detailed biophysical analysis of TCF4 isoform I^-^, which is expressed mainly in the nervous system. This isoform is deficient in amino acids 1-216 in comparison with the canonical isoform B^-^ (UniProt ID P15884-1), and consequently, it lacks the region comprising AD1, the N-terminal NLS and the CE repressor domain, but it contains the entire bHLH domain [4]. Currently, in the literature, there is no information on the molecular properties of TCF4, although the protein has been proposed previously as an integrator (‘hub’) of some of bHLH networks regulating important steps of the CNS developmental program [42]. In fact, some bHLH transcription factors function as transcriptional hubs but their contacts strongly relate to the presence of additional folded motives such as Per-ARNT-Sim (PAS) domains [86]. TCF4 contains no structurally defined domains beside the DNA-binding bHLH domain but still, this protein needs to be able to process various cellular signals and coordinate transcription-related processes. Therefore, it should possess appropriate molecular properties that would allow the protein to form various interactions. The intrinsically disordered character is particularly preferred for proteins with regulatory activity at the crossroads of signaling pathways, as high flexibility allows them to interact with multiple partners in a context-dependent manner [87].

Using the HDX-MS experiment, we demonstrated the fully disordered nature of TCF4 except for the bHLH domain. The disordered conformation is very adaptive, prone to post-translational modifications and often able to fold upon binding [88,89]. The results of the disorder prediction of the TCF4 bHLH domain presented in our previous study [36] suggested the disordered character of the N-terminal, DNA-binding region of bHLH in contrast to the ordered C-terminal part of bHLH. To investigate this phenomenon experimentally, we applied HDX coupled with MS since it is known to reveal the conformation of a polypeptide chain and the conformational changes induced by various factors. They are reflected in changes in isotope exchange [90]. The results of HDX-MS presented in this study indicate that binding of the DNA induces conformational changes resulting only in a greater degree of compactness in the DNA-binding region of TCF4. Folding of the DNA-binding domain upon binding has been observed for the isolated bHLH domain of TCF4 [24] and other TFs [43–45]. Notably, most of the point mutations associated with PTHS occur in the DNA-binding fragment of the bHLH domain [6,29,30,32,91–93]. For this reason, further research focused on this part of the protein may provide a deeper understanding of the molecular basis of TCF4-related diseases. Strikingly, compared with analysis of free TCF4, HDX-MS analysis of TCF4 in complex with the E-box did not reveal any changes in the exchange rate in regions outside the bHLH domain. This finding indicates that binding of the DNA does not induce any conformational change in the regions outside the bHLH domain.

In addition, we demonstrated the ability of TCF4 I^-^ to bind specific DNA with submicromolar affinity. It has been shown that IDRs of some proteins interact with DNA to ease the interaction of a specific DNA sequence with the binding domain [94–96]. Surprisingly, in the TCF4 case, a comparison of the result of the dissociation constant determined for the isolated bHLH domain (K_D_ = 0.11 ± 0.04 μM calculated from the FP for the monomer) [24] versus for the full-length protein (K_D_ = 0.26 ± 0.05 μM calculated from the FP for the monomer) revealed only minor differences.

Notably, dimeric TCF4 I^-^ is one of the largest IDPs expressed and biophysically analyzed to date (Fig. 9) [64,97,98]. Performing shape-related studies of IDPs requires a specific approach [99]. Traditionally, the MW or quaternary structure of globular proteins can be determined on the basis of a direct correlation between the MW and the hydrodynamic properties of the analyzed protein. This principle does not apply to IDPs, which, owing to their dynamic nature, exhibit significantly increased hydrodynamic properties [97]. In this study, we used a less common approach to analyze the conformational changes in disordered proteins induced by ligand binding, i.e., SV-AUC with the f/f_0_ model. A similar approach was previously used by Iconaru et al. [100]. Using this method, we were able to demonstrate that the hydrodynamic properties of dimeric TCF4 in its free and DNA-bound forms are very similar.

**Fig. 9.**
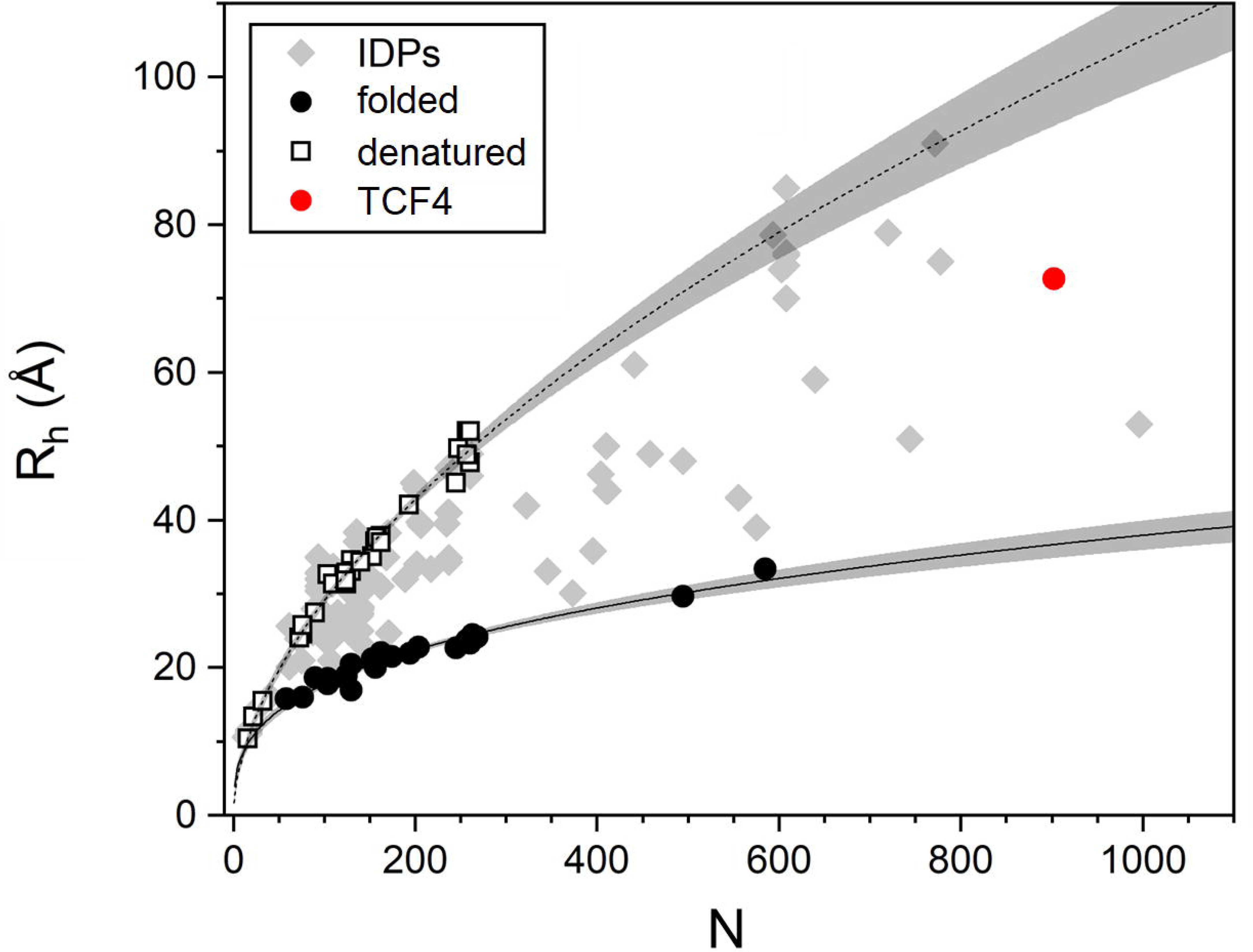
Diagram of the dependence of the hydrodynamic radius on the number of amino acids in the polypeptide chain of known proteins. Gray rhomboids represent IDPs, black circles represent folded proteins, and black empty squares represent denatured proteins described in the literature [64,97,98]. The red circle represents TCF4.

Although specific DNA binding does not affect the disordered nature of TCF4 outside the bHLH domain, we observed that it strongly affects the stability of TCF4. In denaturation experiments, we observed that the native-to-denatured transition of the free form of TCF4 is reversible, but once the TCF4:DNA complex is perturbed, the protein irreversibly loses this property. This suggests that in the DNA-free state, the protein is more flexible, whereas in the *holo* state, it presents lower plasticity. The functional consequences of the altered stability of TCF4 in relation to DNA binding are currently unclear. A flexible structure in the *apo* form clearly has several advantages, i.e., bHLH, which is responsible for both DNA binding and interaction with the dimeric partner, has greater ease of movement. Indeed, the range of motions permitted in an unfolded molecule is less constrained than that permitted in a folded state [101]. This could allow TCF4 molecules in the DNA-free state to dissociate from the wrong dimer partner, providing a second chance to form a complex that can fulfill its function.

We propose that the irreversibility of denaturation of the DNA-bound form of TCF4 could be related to the mechanism by which its activity is inhibited. After transcription initiation, not all the components of the initiation complex dissociate from the promoters [102], and the remaining factors can reinitiate transcription for an extended period of time. Based on our observations indicating that TCF4 is susceptible to irreversible loss of molecular properties only in the DNA-bound state, we propose that certain factors may irreversibly inhibit the activity of some TFs only when they are in the *holo* state. Consequently, TFs can be removed from the pool of active transcription proteins. Notably, the DNA-binding properties of TCF4 and other E proteins are regulated by Ca^2+^-binding proteins such as calmodulin (CaM), S100a and S100b [103–106]. Ca^2+^-dependent proteins interact directly with the basic region of the bHLH domain involved in DNA binding [104], and in the presence of Ca^2+^ can selectively inhibit DNA binding by E proteins [103]. Although it has been shown that altering the levels of CaM or Ca^2+^ in cells has a direct effect on E protein-mediated transcription [107,108], it has not been shown whether CaM binding affects TCF4 stability.

## 7. Conclusions

In conclusion, this study is the first *in vitro* analysis of the molecular properties of the full-length class I bHLH transcription factor. We showed that approximately 90% of TCF4 is composed of disordered regions. Only the HLH domain responsible for dimerization possesses a well-ordered fold. In addition, the basic fragment (b), which is important for E-box binding, actually folds upon binding. These dual structural properties may explain the involvement of TCF4 in the processes that initiate the expression of many genes, allowing TCF4 to act as a “hub” for the regulation of transcription [42]. In this context, we propose that the protein functions in two independent modes. One is related to the function of the bHLH domain and depends on the recognition and binding of the protein to specific DNA sequences, resulting in the anchoring of TCF4 to the appropriate genomic location. Importantly, our observations show that the DNA interaction does not affect the conformation of the fragments outside the binding region. In the other mode, TCF4 IDRs act as a flexible platform for multiple interactions with other transcription-related proteins independent of DNA binding. Although further research is needed to gain a comprehensive understanding of how TCF4 uses the flexibility of its IDRs to participate in the regulation of different biological processes, we believe that our study provides a solid basis for further investigations into the mechanism of the biological activity of TCF4 and other bHLH proteins.

## Supporting information

Supplementary all

## List of abbreviations

*ASCL1:*: Achaete-scute homolog 1

*ATOH1:*: Atonal bHLH transcription factor 1

*bHLH:*: Basic helix-loop-helix

*CaM*: Calmodulin

*CD:*: Circular dichroism

*cv:*: Column volumes

*DOL:*: Degree of labeling

*dsDNA:*: Double-stranded DNA

*dTCF4:*: Dimeric TCF4

*E-box:*: Ephrussi box

*EMSA:*: Electrophoretic mobility shift assay

*FAM:*: 6-carboxy-fluorescein

*FCS:*: Fluorescence correlation spectroscopy

*FP:*: Fluorescence polarization

*HDX-MS:*: Hydrogen/deuterium exchange mass spectrometry

*HES:*: Hairy and enhancer of split

*ID:*: Intellectual disability

*ID-2:*: DNA-binding protein inhibitor ID-2

*IDPs:*: Intrinsically disordered proteins

*IDRs:*: Intrinsically disordered regions

*IMAC:*: Immobilized metal affinity chromatography

*ITF2:*: Immunoglobulin transcription factor 2

*K_D_:*: Dissociation constant

*MW:*: Molecular weight

*PAS:*: Per-ARNT-Sim

*PTHS:*: Pitt-Hopkins syndrome

*R_h_:*: Hydrodynamic radius

*SEC:*: Size exclusion chromatography

*SEF2:*: SL3-3 enhancer factor 2

*SV-AUC:*: Sedimentation-velocity analytical ultracentrifugation

*TCF4:*: Transcription factor 4

*TF:*: Transcription factor

## 8. Declarations

### Ethics approval and consent to participate

Not applicable.

### Consent for publication

Not applicable.

### Availability of data and materials

The datasets generated during and/or analyzed during the current study are available from the corresponding author upon reasonable request.

### Competing interests

The authors declare no competing interests.

### Funding

This work was supported by a subsidy from The Polish Ministry of Science and High Education for the Faculty of Chemistry of Wroclaw University of Science and Technology. The work of B.P.K. and A.N. was partially supported by National Science Centre of Poland Sonata-Bis Grant UMO-2016/22/E/NZ1/00656 to A.N.

### Authors’ contributions

Conceptualization: A.T., B.G.M., A.N., Methodology development: B.P.K., A.N., Investigation: N.S., B.P.L, L.Ż., A.T., Visualization: N.S., B.P.K., A.N., A.T., Writing – Original Draft Preparation: N.S., B.P.K, A.N., B.G.M. A.T., Writing – Review & Editing: N.S., B.P.K., A.N., L.Ż., M.D., B.G.M., A.O., A.T.

## Acknowledgements

We are grateful to Karolina Partyk (WUST) for her excellent technical assistance.

## Authors’ information

Department of Biochemistry, Molecular Biology and Biotechnology, Wrocław University of Science and Technology, Wybrzeże Wyspiańskiego 27, 50-370, Wrocław, Poland

Nikola Sozańska, Beata Greb-Markiewicz, Aneta Tarczewska*, Andrzej Ożyhar

Institute of Physics, Polish Academy of Sciences, Aleja Lotnikow 32/46, 02-668, Warsaw, Poland

Barbara P. Klepka, Anna Niedźwiecka

Institute of Biochemistry and Biophysics, Polish Academy of Sciences, Pawińskiego 5a, 02-106, Warsaw, Poland

Lilia Żukowa, Michał Dadlez

## Notes

### Competing Interest Statement

The authors have declared no competing interest.

