## Supplementary all for "The molecular properties of the bHLH TCF4 protein as an intrinsically disordered hub transcription factor"

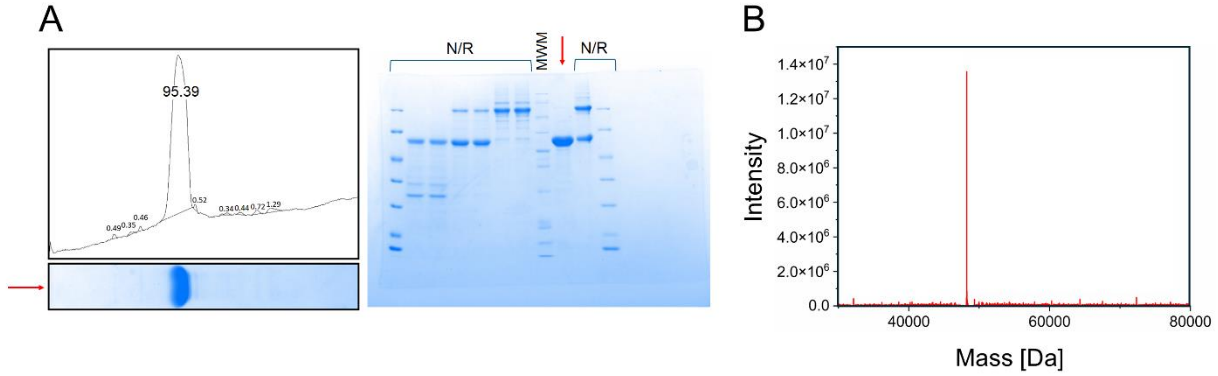


**Fig. S1. Determination of the purity of the TCF4 sample.** (A) Estimation of the purity of the TCF4 sample (red arrow) using ImageJ (left) from SDS-PAGE gel stained with Coomassie Brilliant Blue R-250 (right). The estimated purity of TCF4 was approximately 95%. N/R = not relevant; data that do not contribute to the analysis, because they are unrelated to the context. MWM – Spectra™ Multicolor Broad Range Protein Ladder (Thermo Scientific) (B) TCF4 mass determination via ESI MS. The obtained value equals the theoretical mass of recombinant TCF4 I$^{-}$ (UniProt ID P15884-16) with an additional N-terminal Met.

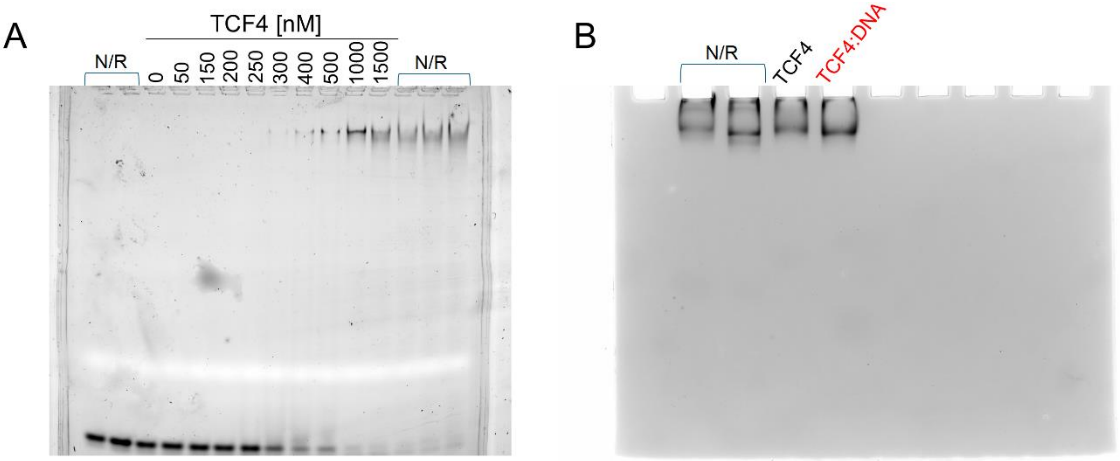


**Fig. S2. TCF4 native PAGE.** (A) E-box binding properties of TCF4 analyzed by EMSA. 10 µl of samples containing a FAM-labeled dsDNA probe (40 nM), TCF4 (0–1.5 µM per dimer), glycerol (final concentration of 20%) and bromophenol blue (final concentration of 1%) were loaded onto 5% native polyacrylamide gels and run at 150 V for 150 min in 0.5 × TBE buffer at 4 °C. Images were captured using the ChemiDoc MP Imaging System (Bio-Rad). (B) Results of TCF4 electrophoresis using 5% native polyacrylamide gel. 20 µl of samples of free TCF4 (2 µM) and TCF4 (2 µM) with E-box (20 µM) were mixed with glycerol (20%) and bromophenol blue (1%). The samples were then loaded onto a gel, and run at 150 V for 40 min in 0.5 × TBE buffer at 4 °C. After electrophoresis, the gel was stained with Coomassie Brilliant Blue R-250 and analyzed using Image Lab Software (Bio-Rad). N/R = not relevant; data that do not contribute to the analysis, because they are unrelated to the context.

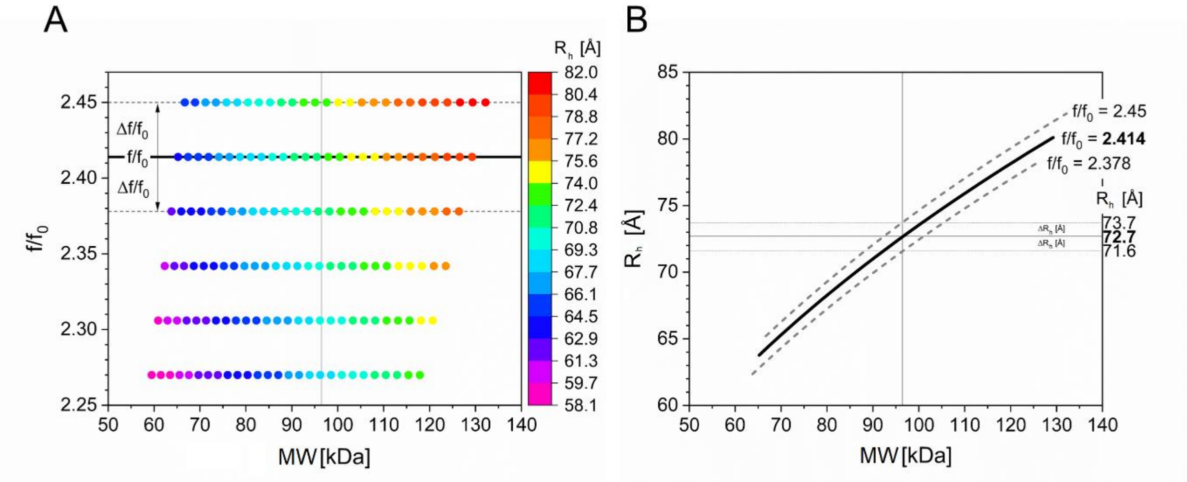


**Fig. S3. The method of determining $f/f_0$ and $R_h$ and their errors using the $c(f/f_0)$ model.** The method assumes a known molecular weight (MW) of the TCF4 dimer. Here, an example is provided for the determination of these values for a 5 µM dTCF4 sample with no DNA ($\bar{\upsilon}$ = 0.7131 mL/g). First, the s distribution for different $f/f_0$ values is calculated. For the s range corresponding to a dimer (A) size ($R_h$) and shape ($f/f_0$) distribution with respect to MW is plotted. The MW of dimer is 96.5 kDa (solid vertical line) and intersects the $f/f_0$ distribution in the middle for $f/f_0$ = 2.414 (solid horizontal line) which is the value that best describes the shape of a dimer. The error of $f/f_0$ ($\Delta f/f_0$) is the difference between $f/f_0$ and the value of the nearest analyzed $f/f_0$ (dashed gray lines). In this case, the $\Delta f/f_0$ is equal to 0.04. Next, (B) $R_h$ as a function of MW is plotted for the determined $f/f_0$ and two adjacent $f/f_0$. The $R_h$ value is the one crossing the $R_h$(MW) function for $f/f_0$ (solid black line) and the $\Delta R_h$ is the difference between $R_h$ and the $R_h$ value determined for the neighboring $f/f_0$ (dashed gray lines).

A

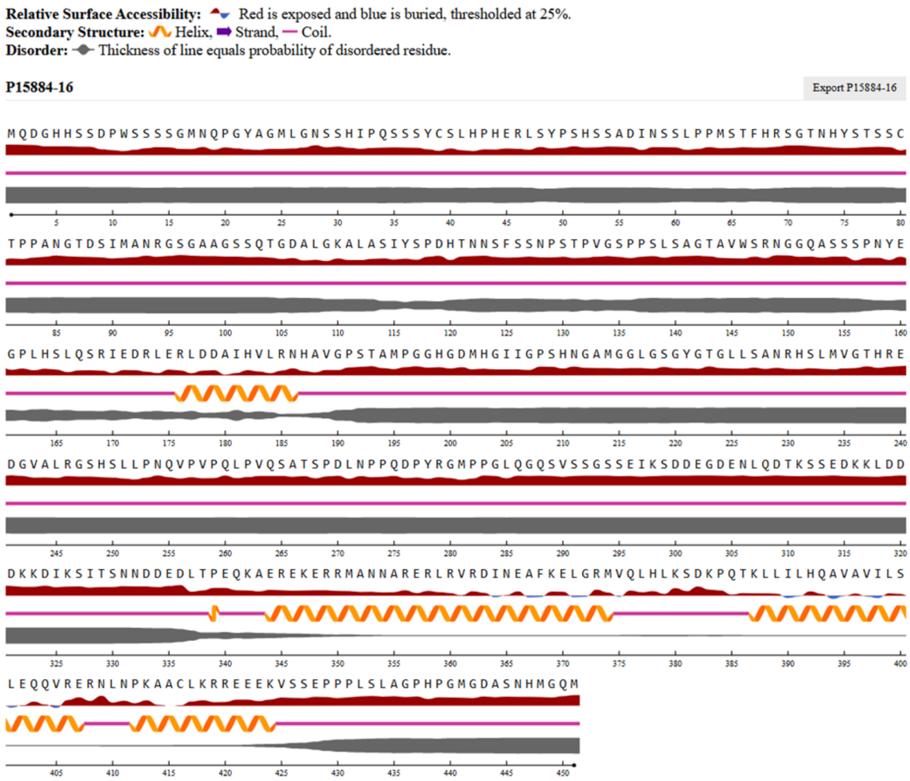


B

| | Helix probability * 10 | | | | | | | | | | | | | | | | | | | | | | | | | | | | | | |
|---|---|---|---|---|---|---|---|---|---|---|---|---|---|---|---|---|---|---|---|---|---|---|---|---|---|---|---|---|---|---|---|
| **No.** | 160 | | | | | | | | | | 170 | | | | | | | | | | 180 | | | | | | | | | | 190 |
| **Residue** | E | G | P | L | H | S | L | Q | S | R | I | E | D | R | L | E | R | L | D | D | A | I | H | V | L | R | N | H | A | V | G |
| **PSIPRED** | 0 | 0 | 5 | 6 | 8 | 9 | 9 | 9 | 9 | 10 | 10 | 10 | 10 | 10 | 10 | 9 | 9 | 10 | 10 | 10 | 10 | 10 | 10 | 10 | 10 | 10 | 9 | 8 | 6 | 2 | 0 |
| **NetSurfP-2.0** | 0 | 0 | 1 | 3 | 3 | 5 | 6 | 6 | 7 | 8 | 9 | 9 | 9 | 10 | 10 | 10 | 10 | 10 | 10 | 10 | 10 | 10 | 10 | 10 | 10 | 10 | 9 | 7 | 4 | 1 | 0 |

**Fig. S4. TCF4 NetSurfP-2.0 and PSIPRED prediction result.** (A) According to the NetSurfP-2.0 results for TCF4, all the residues involved in the formation of secondary structures form the helices. (B) α-helix probability detected for the 166–187 fragment of TCF4 by PSIPRED and NetSurfP-2.0. The consensus predictions are marked in pink.

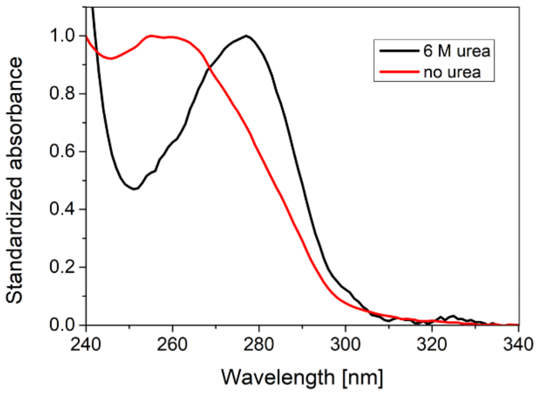


**Fig. S5. UV spectra of TCF4 purified using the 6 M urea rinse step (black) and omitting this step (red).** The apparent shift is due to the presence of nucleic acids in the sample that was not denatured.

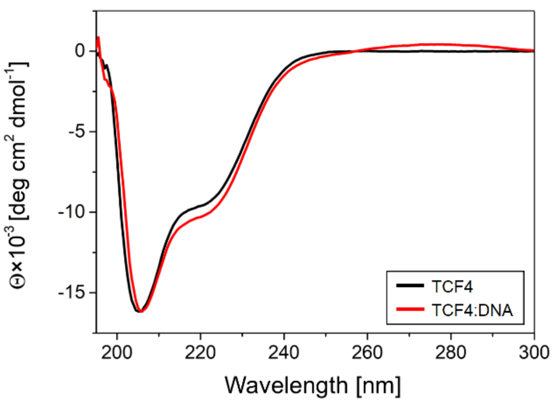


**Fig. S6. CD spectra of TCF4.** TCF4 in both DNA-bound and free form shows a deep minimum around 205 nm and a shoulder at 222 nm typical for a random coil conformation with some contribution of an α-helical structure. In the case of DNA-bound TCF4, the shoulder around 222 nm becomes deeper and the minimum is slightly shifted to 208 nm, suggesting a disorder-to-order transition towards the α-helical fold. A weak signal is also observed at about 280 nm, coming from DNA molecules present in the sample. The analysis using BeStSel deconvolution software revealed that either the free or the DNA-bound form of TCF4 comprise α-helices at approximately 18%.

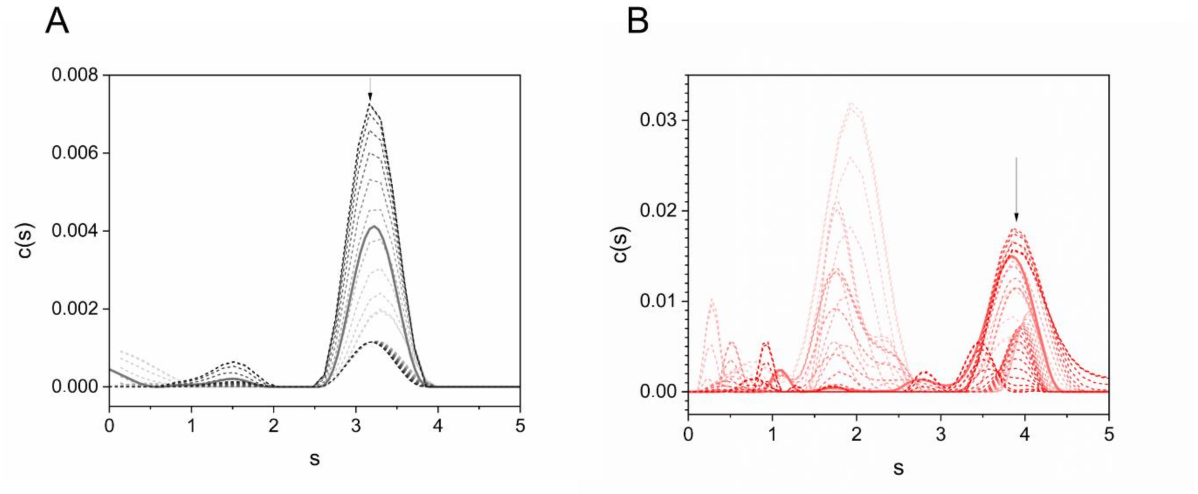


**Fig. S7. Sedimentation coefficient (s) distributions** (dashed and solid lines) of (A) the 5 µM dTCF4 sample with no DNA ($\bar{v}$ = 0.7131 mL/g) and (B) with 10 µM DNA ($\bar{v}$ = 0.697 mL/g) calculated using the c($f/f_0$) model for each analyzed value of $f/f_0$ for different s resolutions. The increasing value of $f/f_0$ was shown as increasing color intensity. The solid lines indicate profiles for (A) $f/f_0$ = 2.41 and (B) $f/f_0$ = 2.32.
